# African army ants at the forefront of virome surveillance in a remote tropical forest

**DOI:** 10.1101/2022.12.13.520061

**Authors:** Matthieu Fritz, Bérénice Reggiardo, Denis Filloux, Lisa Claude, Emmanuel Fernandez, Frédéric Mahé, Simona Kraberger, Joy M. Custer, Pierre Becquart, Telstar Ndong Mebaley, Linda Bohou Kombila, Léadisaelle H. Lenguiya, Larson Boundenga, Illich M. Mombo, Gael D. Maganga, Fabien R. Niama, Jean-Sylvain Koumba, Mylène Ogliastro, Michel Yvon, Darren P. Martin, Stéphane Blanc, Arvind Varsani, Eric Leroy, Philippe Roumagnac

**Affiliations:** Institut de Recherche pour le Développement (IRD), Maladies Infectieuses et Vecteurs, Ecologie, Génétique, Evolution et Contrôle (MIVEGEC) (Université de Montpellier-IRD 224–CNRS 5290), 34394 Montpellier, France; CIRAD, UMR PHIM, 34090 Montpellier, France; PHIM Plant Health Institute, Univ Montpellier, CIRAD, INRAE, Institut Agro, IRD, Montpellier, France; The Biodesign Center for Fundamental and Applied Microbiomics, Center for Evolution and Medicine, School of Life Sciences, Arizona State University, Tempe, AZ, USA; Centre Interdisciplinaire de Recherches Médicales de Franceville, Franceville, Gabon; Laboratoire National de Santé Publique, Brazzaville, Republic of Congo; Université Marien Ngouabi, Brazzaville, Republic of Congo; Department of Anthropology, Durham University, South Road, Durham DH1 3LE, UK; Université des Sciences et Technique de Masuku (USTM), Institut National Supérieur d’Agronomie et de Biotechnologies (INSAB), Franceville, Gabon; INRA-Université de Montpellier UMR DGIMI 34095 Montpellier, France; Division of Computational Biology, Department of Integrative Biomedical Sciences, Institute of infectious Diseases and Molecular Medicine, University of Cape Town, Cape Town, South Africa; Structural Biology Research Unit, Department of Integrative Biomedical Sciences, University of Cape Town, Observatory, Cape Town, South Africa

**Author notes:** These authors contributed equally to this work (Matthieu Fritz, Bérénice Reggiardo, Eric Leroy, Philippe Roumagnac).

## Abstract

In this study, we used a predator-enabled metagenomics strategy to sample the virome of a remote and difficult-to-access densely forested African tropical region. Specifically, we focused our study on the use of army ants of the genus *Dorylus* that are obligate collective foragers and group predators that attack and overwhelm a broad array of animal prey. Using 209 army ant samples collected from 29 colonies and the virion-associated nucleic acid-based metagenomics approach, we showed that a broad diversity of bacterial, plant, invertebrate and vertebrate viral sequences were accumulated by army ants: including sequences from 157 different viral genera in 56 viral families. This suggests that using predators and scavengers such as army ants to sample broad swathes of tropical forest viromes can shed light on the composition and the structure of viral populations of these complex and inaccessible ecosystems.

## Introduction

Viruses are likely the most abundant and diverse biological entities on Earth ^1–3^ and are arguably the most successful inhabitants of the biosphere ^4^. Despite this, the current inventory of known virus diversity is likely a vanishingly small and unrepresentative fraction of the total diversity, abundance, and population structures of all extant viruses ^1, 5–7^. Hence, while the overall total number of eukaryotic virus species on Earth is estimated to be in excess of several million ^1^, only 10,434 virus species (including eukaryotic and prokaryotic viruses) are presently recognized by the International Committee on Taxonomy of Viruses ^8^. Our perception of the true extent and properties of the virosphere is further clouded by the fact that this minuscule sample of global virus diversity is heavily biased towards virus species that directly impact humans and the organisms that we tame and farm ^6, 7^.

The best studied components of the virosphere are those that include the plant, animal, fungi and bacteria-infecting viral agents – called viromes – within or upon human bodies, human food sources (especially domesticated plants and animals) and human habitats (especially urban homes, hospitals, schools and farms). Conversely, the least studied components of the virosphere are those of natural environments, particularly remote ecosystems such as those found in deep tropical forests. Crucially, even when metagenomic projects have explored the viromes in such regions, these studies have been geographically and taxonomically biased. Hence, samples were only accessible via forest roads or tracks, and derived from the subset of plant or animal species that are likely to host viruses with some medical or agricultural relevance ^9^. Such sampling biases are understandable in that most future viral diseases of humans and their domesticated plants and animals will likely emerge from host species living or wandering in their vicinity. In addition, future emerging diseases will often be related to viruses already known to cause diseases in humans, and domesticated plants and animals. However, it is important to keep in mind that a less human-centric assessment of viral diversity at the ecosystem-scale could i) illuminate the natural host-ranges, ecological contexts and evolutionary processes underlying the diversification of viruses related to those that have already emerged to cause diseases that impact humans, and ii) identify the yet unknown viruses with properties such as broad host ranges or high incidences that could potentially pose future threats to humans ^9, 10^. More generally, the paucity of information on virus diversity in natural environments is hampering our understanding of both the roles of viruses within wild ecosystems, and how natural- or human- mediated disturbances of such ecosystems might impact these roles ^10–12^.

Densely forested tropical regions account for 40 % of the world’s 4 billion hectares of forests ^13^ and provide, as a consequence of human activities surrounding these forests, major interfaces where humans interact with the world’s remaining wilderness areas. Besides accessibility issues, studying viromes in these habitats poses several additional challenges. The spatial pattern of sampling schemes, the establishment of a list of organisms to sample, and the developmental stages and/or symptom statuses of these organisms are all important considerations as they may or may not yield metagenomic data representative of the diversity of viruses circulating within the targeted area ^14^. An interesting “meta-sampling” strategy that provides an alternative to classical human-centric assessment of viral diversity in inaccessible sampling sites is to rely on proxy samplers such as highly mobile predator/scavenger animals that naturally accumulate animal-, fungal- and plant-derived biomass within their digestive tracts during feeding ^15–19^. Rather than aiming for completely random sampling of biologicalmaterial within a given environment, the general intention of this approach, -which is variously referred to as “xenosurveillance” or “vector-enabled metagenomics” (VEM) - is to shift the causes of unrepresentative sampling from human choice biases to the feeding-choice biases of the chosen mobile scavenger/predator that is intimately involved with the targeted environments ^15–19^. While feeding-choice biases are unavoidable with such meta-sampling schemes, they are biologically meaningful in that viral cargos of blood-feeding, sap-feeding and insectivorous insects are themselves inherently representative of viral mobility within ecosystems. Insectivorous arthropods, so-called top-end insect predators, are potentially particularly useful samplers in this regard. Indeed, the diversity of viruses that they ingest via their prey should be partly representative of the viruses that are ferried within insects throughout the ecosystem. In addition, sampling top-end insect predators will avoid the capture (and sometimes sacrifice) of animals that are protected for ethical, legal or cultural reasons. Among xenosurveillance approaches, Predator Enabled Metagenomics (PEM), using for example dragonflies^20^ or damselflies, ^21^ has proved more efficient with respect to collecting a wide diversity of viral sequences than Vector-Enabled Metagenomics (VEM) using exclusively plant sap-feeding insects (e.g. whiteflies ^19^) or blood-feeding insects (e.g. mosquitoes ^15–17^).

In this study, we propose a PEM strategy using army ants of the genus *Dorylus* ^22, 23^ as top-end insect predators. These nomadic insects are obligate collective foragers and group predators that attack a broad array of animal prey including crickets, cockroaches, earthworms, and even vertebrates. Although nomadic, they do have temporary nests around which they daily forage areas of ∼1km ^2, 22–24^. Army ants generally live in colonies containing between 10^4^ to 10^7^ workers ^25^ with a daily colony-wide intake of prey/scavenged biomass of up to 2 kg ^26^. In addition to preying on an extremely diverse array of animal species, army ants also scavenge on the carcasses of large vertebrates and feed directly on plants ^24, 27, 28^.

Here we test the overarching hypothesis that army ants hunting live invertebrate and vertebrate prey in the deep forest ingest and accumulate a diverse array of plant and animal viruses in the areas around their temporary nests.

## Materials and Methods

### Army ants sampling

In July 2019, two sampling surveys were conducted in the Ogooué-Ivindo region (Northeast Gabon). Over 250 individual ants from 29 colonies were collected along roadsides between Mekambo and Mendemba villages, immediately stored in liquid nitrogen and further transferred to Montpellier (France) where they were stored at -80°C. Two-hundred and nine samples, including 145 samples containing only one ant and 64 samples each containing pools of 2 to 13 individual ants (Table 1 and Supplementary Table 1) were further processed using the virion-associated nucleic acid based (VANA) metagenomics approach ^29^.

**Table 1:** Characteristics of the army ant samples. P21/P23 means that colonies P21 and P23 were pooled. The species assignation is based on Cytochrome oxidase I reads and contigs analysis. “nd” means non determined.

| Ant colony number | Nb. of samples | Genus | Species | Foraging niche | Collection date | Collection location | GPS coordinates |
| --- | --- | --- | --- | --- | --- | --- | --- |
| FM1 | 4 | *Dorylus* sp. | *D. sjoestedti/wilverthi* clade | surface swarms | 2019-07-07 | Mendemba | nd |
| FM2 | 4 | *Dorylus* sp. | *D. sjoestedti/wilverthi* clade | surface swarms | 2019-07-07 | Mendemba | nd |
| FM3 | 4 | *Dorylus* sp. | nd | nd | 2019-07-07 | Between Ntolo and Mekouma | nd |
| FM4 | 4 | *Dorylus* sp. | *D. sjoestedti/wilverthi* clade | surface swarms | 2019-07-07 | Between Ntolo and Mekouma | nd |
| FM5 | 4 | *Dorylus* sp. | *D. sjoestedti/wilverthi* clade | surface swarms | 2019-07-07 | Mekouma | nd |
| FM6 | 4 | *Dorylus* sp. | nd | nd | 2019-07-07 | Between Mekouma and Malassa | nd |
| FM7 | 4 | *Dorylus* sp. | *D. sjoestedti/wilverthi* clade | surface swarms | 2019-07-07 | Mekambo | nd |
| FM8 | 4 | *Dorylus* sp. | nd | nd | 2019-07-07 | Mekambo | nd |
| P1A | 9 | *Dorylus* sp. | nd | nd | 2019-07-08 | Mekambo | 00°57.214' - 013°58.396' |
| P2A | 5 | *Dorylus* sp. | *D. sjoestedti/wilverthi* clade | surface swarms | 2019-07-08 | Komambela | 00°49.865' - 013°58.846' |
| P3A | 8 | *Dorylus* sp. | *D. sjoestedti/wilverthi* clade | surface swarms | 2019-07-09 | Mekambo | 00°56.681' - 013°59.032' |
| P4A | 8 | *Dorylus* sp. | *D. sjoestedti/wilverthi* clade | surface swarms | 2019-07-09 | Between Ibong and Zoula | 01°02.872' - 014°00.951' |
| P5 | 8 | *Dorylus* sp. | *D. sjoestedti/wilverthi* clade | surface swarms | 2019-07-09 | Between Ibea and Zoula | 01°04.409' - 014°03.100' |
| P6 | 8 | *Dorylus* sp. | *D. opacus/kohli* clade | leaf-litter | 2019-07-09 | Between Zoula and Grand Etoumbi | 01°08.545' - 014°06.826' |
| P7 | 8 | *Dorylus* sp. | *D. sjoestedti/wilverthi* clade | surface swarms | 2019-07-09 | Between Massombo and Egopouma | 01°16.394' - 014°06.873' |
| P8A | 9 | *Dorylus* sp. | nd | nd | 2019-07-09 | Between Ibea and Zoula | 01°04.086' - 014°02.629' |
| P9 | 9 | *Dorylus* sp. | nd | nd | 2019-07-09 | Between Ibea and Zoula | 01°03.571' - 014°02.068' |
| P10 | 9 | *Dorylus* sp. | nd | nd | 2019-07-10 | Between Mbessa and Malassa | 00°54.502' - 014°01.582' |
| P11 | 9 | *Dorylus* sp. | nd | nd | 2019-07-12 | Between Malassa and Mekouma | 00°51.149' - 014°04.198' |
| P12 | 9 | *Dorylus* sp. | *D. sjoestedti/wilverthi* clade | surface swarms | 2019-07-12 | Between Ntolo and Mekouma | 00°49.975' - 014°04.212' |
| P13 | 9 | *Dorylus* sp. | nd | nd | 2019-07-10 | Between Mekouma and Ntolo | 00°48.731' - 014°04.110' |
| P14 | 9 | *Dorylus* sp. | *D. sjoestedti/wilverthi* clade | surface swarms | 2019-07-10 | Between Mekouma and Ntolo | 00°44.720' - 014°05.251' |
| P15 | 9 | *Dorylus* sp. | *D. sjoestedti/wilverthi* clade | surface swarms | 2019-07-11 | Mekambo | 00°59.282' - 013°57.916' |
| P16 | 9 | *Dorylus* sp. | *D. sjoestedti/wilverthi* clade | surface swarms | 2019-07-11 | Mekambo | 00°58.182' - 013°58.277' |
| P17 | 9 | *Dorylus* sp. | *D. sjoestedti/wilverthi* clade | surface swarms | 2019-07-11 | Mekambo | 00°57.042' - 013°58.532' |
| P18 | 9 | *Dorylus* sp. | nd | nd | 2019-07-11 | Mbeza | 00°55.154' - 013°59.798' |
| P19 | 9 | *Dorylus* sp. | nd | nd | 2019-07-12 | Between Mekambo and Mbeza | 00°56.939' - 013°58.605' |
| P21/P23 | 5 | *Dorylus* sp. | *D. sjoestedti/wilverthi* clade | surface swarms | 2019-07-12 | Mbeza | 00°55.167' - 013°59.793' / 00°52.804' - 014°03.581' |
| P24 | 10 | *Dorylus* sp. | nd | nd | 2019-07-12 | Between Malassa and Mekouma | 00°52.253' - 014°03.609' |

### Virion-associated nucleic acid-based viral metagenomics

Each of the 209 collected samples was processed using the VANA viral metagenomics approach that is comprehensively detailed in François et al. (2018) ^30^. It is noteworthy that the VANA approach is suited to the detection of both DNA and RNA viruses, and includes several steps that aim at both removing host nucleic acids and maximizing the yield of virion-associated nucleic acids ^29^. Briefly, ants were first ground using tissue homogenizer and sterile steel beads. Viral particles from individual or pooled ant samples were first isolated using centrifugation and filtration techniques and they were further concentrated by ultracentrifugation. Contaminating non-encapsidated nucleic acids were then digested by DNase and RNase digestion treatments. Following this, encapsidated DNA and RNA molecules resistant to the DNase and RNase treatments were extracted. A series of molecular amplification was subsequently carried out, including reverse transcription, Klenow fragment treatment, and amplification of the viral DNA and RNA using barcoded PCR primers. Finally, amplification products were pooled into 3 libraries. Five negative controls, each containing 8 ml of 1x Hanks’ buffered salt solution, were also added to the three libraries. The ant samples and the negative controls were further sequenced by Genewiz (Leipzig, Germany) using a single lane on an Illumina HiSeq 3000/HiSeq 4000 sequencer (2 × 150 bp sequencing). Bioinformatics analyses were performed as described previously ^30^. Briefly, demultiplexing was performed with the agrep command-line tool to assign reads to the samples from which they originated ^31^. Adaptors were removed and the reads were filtered for quality (q30 quality and read length >45 nt) using Cutadapt 3.1 ^32^. The cleaned reads were assembled *de novo* into contigs using SPAdes 3.6.2 ^33^. Putative virus reads obtained using BLASTx ^34^ against the GenBank non-redundant protein database with e-values < 0.001 were retained. Amplification products (amplicons) of four samples (N° 166, 185, P13-3.2 and P17-2.2, Supplementary Table 1) were also sequenced in parallel using the recently developed Flongle (flow cell dongle) sequencing system (Oxford Nanopore Technologies, Oxford, UK). These amplicons were purified using Agencourt AMPure XP beads (Beckman Coulter, Brea, CA, USA). Library construction for the Flongle sequencing system was performed using the SQK-LSK109 Kit, following the manufacturer’s instructions. Four Flongle Flow Cell (R9.4.1) were used for sequencing. The bioinformatics analysis of the Nanopore reads was carried out as follows: accurate base calling was performed using Guppy (v5.0.16; available online at https://nanoporetech.com/). Adapter and primer (Dodeca linker) removal was then performed using Porechop v0.2.4 (available online at https://github.com/rrwick/Porechop). The quality of reads was investigated using NanoPlot v1.33.0 ^35^. Taxonomic assignment was achieved on cleaned Nanopore reads through searches against the NCBI nr protein database using DIAMOND 0.9.22 with an e-value threshold of < 0.001 ^36^. Read analyses of the five negative controls were performed and numbers of virus reads were determined. Indicative of cross-sample contamination, virus reads assigned to the virus families mostly represented in the ant samples (i.e. *Parvoviridae*, *Microviridae*, *Dicistroviridae*, *Circoviridae*, *Iflaviridae*, *Polycipiviridae*, *Retroviridae*, *Bidnaviridae* and *Nodaviridae*) as well as cruciviruses were found associated with the negative control, with a mean of 15 reads per virus family. This result suggests that a minimum of 15 reads assigned to these 10 virus families would be a conservative threshold above which a sample should be considered as likely containing viral sequence assigned to these families. On the other hand, no “read threshold” was used for the virus families for which no evidence of cross-sample contamination was identified.

### Dorylus sp. cytochrome oxidase I reads inventory and taxonomic assignment

Illumina reads assigned to *Dorylus* sp. *cytochrome oxidase I* gene using BLASTx searches were recovered and further assembled using SPAdes ^33^. Contigs and representative *cytochrome oxidase I* gene sequences of *Dorylus* sp. specimens representing all six recognized subgenera ^37^ were subsequently aligned using MUSCLE with default settings ^38^. A phylogenetic tree was constructed using the maximum likelihood method implemented in PhyML 3.1 ^39^. The HKY85 substitution model was selected assuming an estimated proportion of invariant sites of 0.542 and 4 gamma-distributed rate categories to account for rate heterogeneity across sites. The gamma shape parameter was estimated directly from the data (gamma=1.219). Support for internal branches was assessed using the aLRT test (SH-Like).

### Inventory of virus contigs and putative taxonomic assignment

One of the critical aspects of using BLAST-based searches to assign contigs to viral families is the minimum length of the contig being queried. We have recently conducted a simulation experiment that has revealed that the accuracy of the BLASTx virus family and genus assignations are high, *i.e.* >97.9% and >90.6%, respectively, when contigs have lengths ≥200 nt ^29^. We therefore selected contigs with lengths ≥200 nt and retained viral BLASTx assignations of these contigs wherever they yielded e-values < 0.001.

### Statistical analyses

The taxonomic assignations of ≥200 nt long virus contigs at the genus level (as determined by best BLASTx hits) were scored for individual ants, including 67 individual workers and 78 individual soldiers. Differences between the number of viral genera associated with workers and soldiers were compared using Mann-Whitney U-tests. Differences between the number of viral genera associated with individual ants belonging to the *Dorylus sjoestedti/wilverthi* clade (n = 76) and to the *Dorylus opacus/kohli* clade (n = 6) were also compared using Mann-Whitney U-tests. Finally, differences between the number of viral genera associated with individual ants from 20 *Dorylus* colonies (4 ≤ n ≤ 8) were compared using the Kruskal-Wallis H test.

### Phylogenetic analyses of most prevalent virus families

Contigs assigned to the families Bidnaviridae, Dicistroviridae, Hepeviridae, Iflaviridae, Microviridae, Nodaviridae, Picobirnaviridae, Polycipiviridae, Tombusviridae, Tymoviridae and Solemoviridae, as well the Picorna-like virus group and the CP-based sequence grouping of viruses called cruciviruses ^40^, were translated and conserved protein sequences were extracted (the even more prevalent parvo- and cycloviruses are analysed separately, see next sections). These protein sequences together with representative sets of protein sequences belonging to the virus groups/families to which the sequences were taxonomically assigned, were aligned using MUSCLE with default settings ^38^. Neighbor joining phylogenetic trees were generated using MEGA version X ^41^ using alignments of major capsid protein sequences (Bidnaviridae and Microviridae), capsid protein sequences (CP; cruciviruses), RNA-dependent RNA polymerase sequences (RdRp; Dicistroviridae, Nodaviridae, Picobirnaviridae and Solemoviridae), polyprotein sequences (Hepeviridae, Iflaviridae, Picorna-like viruses, Tombusviridae and Tymoviridae), and ORF5 sequences (Polycipiviridae). One thousand bootstrap replicates were performed to quantify branch support.

### Phylogenetic analyses of parvovirus-related sequences

We attempted to evaluate the genetic relationships of the parvovirus-related sequences (PRS) that were by far the most abundant virus sequences amplified from the army ant samples. To reconstruct the evolutionary relationships of the various major parvovirus lineages we focused exclusively on the SF3 (Super Family 3) helicase domain of parvovirus protein, NS1. The SF3 helicase domain is highly conserved in all known parvoviruses and is therefore typically used for phylogenetic analyses of divergent parvoviruses ^42, 43^. NS1 BLASTx assignations (with e-values < 0.001) were initially retained and translated *in silico* using ORF finder (cut off ≥ 500 bp) (http://www.ncbi.nlm.nih.gov/projects/gorf/). NS1 protein sequences that were ≥200 aa in length were then selected and processed using the software Interproscan which predicted the presence of functional domains ^44^. Four-hundred and three SF3 protein sequences were further selected and combined with both 125 representative SF3 protein sequences of viruses from publicly available transcript, genome and protein databases that were classified as belonging to the *Parvoviridae* family ^43^ and 155 SF3 protein sequences collected from available genomes of viruses classified in the *Parvoviridae* family from the NCBI genome database. The 683 SF3 protein sequences were aligned using MUSCLE with default settings ^38^. A Neighbor-Joining tree was produced using MEGA version X ^41^ with 1000 bootstrap replicates to quantify branch support. Specifically, NS1 protein sequences that were ≥200 aa in length and which were assigned to the genus *Chaphamaparvovirus* were aligned together with 32 representative protein sequences from this genus using MUSCLE with default settings ^38^. A phylogenetic tree was constructed using the maximum likelihood method implemented in PhyML 3.1 ^39^. The WAG substitution model was selected by PhyML 3.1 assuming an estimated proportion of invariant sites of 0.075 and four gamma-distributed rate categories to account for rate heterogeneity across sites. The gamma shape parameter was estimated directly from the data (gamma=1.337). Support for internal branches was assessed using the aLRT test (SH-Like).

### Sequencing and phylogenetic analyses of complete cyclovirus genomes

Four hundred and seventy-two contigs ≥ 200 nt in length that shared similarity with members of the *Circoviridae* family were identified using BLASTx searches with e-values < 0.001 (Supplementary Table 2). These contigs were initially aligned using MUSCLE and clustered in 22 genetic groups (data not shown). Abutting primer pairs were designed that were specific to these genetic groups (Supplementary Table 3) to enable the recovery of full viral genomes using polymerase chain reaction (PCR) with HiFi HotStart DNA polymerase (Kapa Biosystems, USA) using cycling conditions per the manufacturer’s instructions. The amplicons were resolved on a 0.7% agarose gel using electrophoresis, and amplicons approximately 2–3 kb in size were excised from the gel and purified using the Quick-spin PCR Product Purification Kit (iNtRON Biotechnology, Korea). The amplicons were further cloned in pJET1.2 cloning plasmid (ThermoFisher Scientific, USA), and the recombinant plasmids were then transformed into XL1 blue *Escherichia coli* competent cells. The resulting plasmids from the transformants were purified using a DNA-spin Plasmid DNA Purification kit (iNtRON Biotechnology, Korea), and then sequenced by primer walking at Macrogen Inc. (Korea). Specifically, 45 translated Rep sequences recovered from 45 complete genome sequences and representative cyclovirus Rep protein sequences were aligned using MUSCLE with default settings ^38^. A phylogenetic tree was constructed using the maximum likelihood method implemented in PhyML 3.1 ^39^. The LG+I+G substitution model was selected assuming an estimated proportion of invariant sites of 0.075 and 4 gamma-distributed rate categories to account for rate heterogeneity across sites. The gamma shape parameter was estimated directly from the data (gamma=1.337). Support for internal branches was assessed using the aLRT test (SH-Like).

## Results and discussion

### Classification of army ants using mitochondrial cytochrome oxidase I gene

Before examining the diversity of viruses associated with the army ant samples that we collected from Gabon, we attempted to determine the *Dorylus* species that the ants belonged to by examining their mitochondrial *cytochrome oxidase I* genes. Ninety-nine reads from 24/209 of the ant samples, representing 17/29 of the ant colonies, that shared identity with the *cytochrome oxidase I* gene were recovered from the Illumina sequencing run. Sixty-four of these 99 reads were further assembled into five contigs with lengths ranging from 123 nt to 225 nt. Two maximum likelihood phylogenetic trees containing these contigs together with partial sequences of the *cytochrome oxidase I* gene of 38 *Dorylus* sp. specimens (representing all six recognized *Dorylus* subgenera) indicated that four contigs (contigs #1, #3, #4 and #5) clustered with *Dorylus sjoestedti*, *Dorylus wilverthi* and *Dorylus rubellus* whereas the last contig (contig #2) clustered with *Dorylus opacus*, *Dorylus kohli* and *Dorylus helvolus* (Supplementary Figure 1). While contig #2 comprised seven reads obtained from two samples of colony P6, contigs #1, #3, #4 and #5 were composed of 57 reads obtained from 23 samples from 15 colonies (Table 1 and Supplementary Table 1).

In addition, 266 reads that shared identity with the *cytochrome oxidase I* gene were recovered from four Nanopore sequencing runs. These Nanopore reads shared high identity with *Dorylus sjoestedti* and *Dorylus wilverthi* suggesting that ant samples from the FM1 (not assigned by Illumina reads), FM2 and P17 colonies were also related to these two army ant species. Finally, no *cytochrome oxidase I* reads were identified from eleven ant colonies that, therefore, remained taxonomically unassigned (Supplementary Table 1).

*Dorylus sjoestedti*, *Dorylus wilverthi* and *Dorylus rubellus* are army ant species that are considered “swarm foragers” (most commonly known as driver ants) on the forest floor and in the lower vegetation ^37^. In contrast, *Dorylus opacus* and *Dorylus kohli* which hunt small invertebrates and worms in the leaf-litter are more closely related to subterranean species of *Dorylus* sp., than to the surface swarm foraging ants ^37^. Overall, the mitochondrial *cytochrome oxidase I* gene analysis confirmed that the army ants collected in our study were mostly of the genus *Dorylus* and that they were therefore suitable for testing our overarching hypothesis that army ants hunting live invertebrate and vertebrate prey in the deep forest ingest and accumulate numerous plant and animal viruses present in an area around their temporary nests.

### Genetic and morphological factors influence the army ant virome

The three sequencing libraries each contained an average of 50 million reads following removal of short (<15 nt) sequences and individual sequence regions with low quality scores. Overall, 443,645 contigs that were ≥200 nt in length were assembled and 46,377 of these contigs (10.5%) exhibited sequence similarity to viruses (BLASTx e-values < 0.001). Among these 46,377 contigs, 11,146 were assigned at the viral realm level, 1,377 were assigned at the viral order level and 11,448 were assigned at the viral family level. Only 22,406 of the 46,377 contigs (48.3%) exhibited sequence similarity to viral genera recognized or in the process of being recognized by the International Committee on Taxonomy of Viruses. These apparently virus-derived contigs yielded detectable homology to viruses of 157 different viral genera in 56 viral families (Supplementary Table 2); overall, a higher diversity than the 29 virus families identified from queens from 49 colonies of North American red fire ant *Solenopsis invicta* ^45^. It is noteworthy that the most frequently amplified viral sequences belonged to families containing viruses that are known to infect invertebrates, including *Circoviridae*, *Dicistroviridae*, *Iflaviridae*, *Polycipiviridae* and *Parvoviridae* (Figure 1, Supplementary Table 2). However, plant-, bacteria- and vertebrate-associated viruses were also detected (Figure 1, Supplementary Table 2). While the diversity of the detected virus genera was apparently important, several virus genera that are ubiquitous worldwide (such as *Amalgavirus*, *Endornavirus*) were either undetected or rarely detected. This may be attributable to the metagenomics approach that was used being based on the analysis of virion-associated nucleic acids: an approach that is potentially biased towards the semi-purification of viral capsids that are resilient to harsh conditions within the digestive tracts of the ants and the organisms that they prey on. This may explain why capsidless RNA viruses of fungi, oomycetes and plants, such as those in the families *Amalgaviridae* (not detected), *Deltaflexiviridae* (1 contig detected), *Endornaviridae* (2 contigs detected), *Hypoviridae* (no contigs detected), *Mitoviridae* (no contigs detected), *Narnaviridae* (1 contig detected) and *Polymycoviridae* (no contigs detcetd) ^46, 47^ were either undetected or only rarely detected in this study.

**Figure 1:**
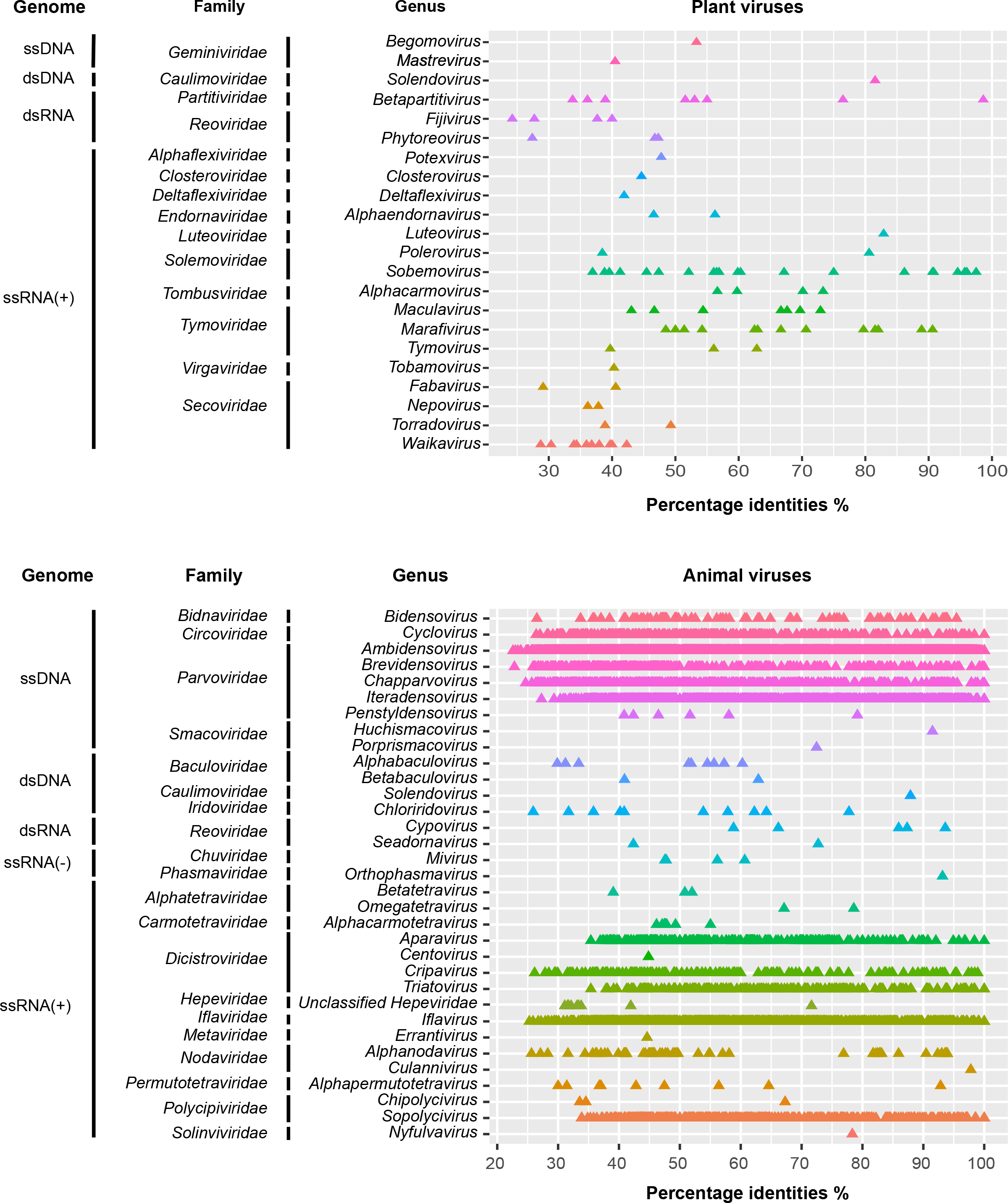
Sequence identity distribution analysis of viral sequences from the army ant virome. Each dot represents an assembled sequence contig with the corresponding protein identity (best BLASTx e-values < 0.001) to plant (top) and animal virus (bottom) in the GenBank nonredundant database.

The taxonomic assignations of virus contigs ≥200 nt in length at the genus level were scored for individual ants belonging to 20 *Dorylus* colonies, including 67 individual workers and 78 individual soldiers. No significant difference between the number of virus genera identified from individual ants of the 20 *Dorylus* colonies were found (p-value = 0.1067), suggesting that our roadside sampling survey in the Ogooué-Ivindo region of Gabon was reasonably homogeneous. Remarkably, ants belonging to the *Dorylus opacus/kohli* clade yielded contigs with homology to viruses in significantly fewer genera (median = 4.5, SD = 2.32) than ants assigned to the *Dorylus sjoestedti/wilverthi* clade (median = 9, SD = 4.02; p-value = 0.0057).

While ant species in the *Dorylus opacus/kohli* clade hunt in leaf-litter and forage on forest floors and on vegetation, the driver ant species from the *Dorylus sjoestedti/wilverthi* clade are swarm raiders ^25^ that prey on invertebrates and occasionally on vertebrates ^37^. These driver ants therefore potentially hunt a wider diversity of prey that may result in the intake of a larger diversity of viruses. This result needs to be further confirmed, however, since only eight ants from the *Dorylus opacus/kohli* clade could be analyzed in this study (Table 1).

Finally, we found that contigs obtained from worker ants that were homologous to known viruses represented significantly fewer virus genera (median = 7, SD = 3.52) than those obtained from soldier ants (median = 10, SD = 4.27; p-value = 0.00007). Whereas worker ants specialize in hunting and collecting food, soldier ants have powerful mandibles and specialize in colony defence. In addition, the workers have smaller body sizes than soldier ants. At this stage, we have no clear explanation as to why the diversity of viruses was higher in the soldier ants than in the worker ants. We can hypothesise that i) soldiers simply eat more than workers, soldiers ingest a greater diversity of organisms than workers because some of the animals that are attacked during defence are not necessarily food for the colony, and/or iii) have an immune system that is more permissive of viral infections. Of relevance is also the fact that host body size frequently shows a positive correlation with parasite species richness ^48, 49^, large-bodied soldier ants may provide greater cellular capacity for viral storage/replication than small-body worker ants. Whatever the explanation, this observation suggests that driver-ant soldiers may be ideal candidates for the surveillance of viruses circulating in tropical forest ecosystems.

### A broad diversity of bacterial, plant, invertebrate and vertebrate viruses is accumulated by army ants

#### Bacteriophages

Overall, 583 contigs ≥200 nt in length (mean size = 286 nt) were assigned to five bacteriophage families (*Herelleviridae*, *Microviridae*, *Namaviridae*, *Picobirnaviridae* and *Tectiviridae*) and three “unclassified Caudoviricetes” (recently abolished *Myoviridae*, *Podoviridae* and *Siphoviridae* families ^8^) with *Microviridae* accounting for 83.7% (488) of these: a result suggesting that the VANA approach may not efficiently detect bacteriophages with a head and tail structure (Supplementary Table 2). Phylogenetic analyses of the major capsid protein sequences of 30 contigs assigned to the *Microviridae* revealed that these could be subdivided into three groups (Figure 2): (1) one comprising 21 contigs branching with an unclassified *Gokushovirinae* isolate (isolate Bog1183 53; accession number: YP_009160331) recovered from a sphagnum-peat soil ^50^; (2) one comprising four contigs clustering with Spiroplasma virus SpV4; Acc. Nbr: NP_598320); and (3) one comprising five contigs grouping with two unclassified microviruses (Alces alces faeces associated microvirus MP21 4718 isolated from moose feces [Acc. Nbr: YP_009551424] and Fen7940 21 [Acc. Nbr: YP_009160412] isolated from a sphagnum-peat soil). Additionally, 33 contigs were assigned to the *Picobirnaviridae* family (Supplementary Table that comprises viruses putatively infecting prokaryotes ^51^. Our phylogenetic analyses showed that three army ant contigs (for which we obtained RdRp sequences >130 aa) were unambiguously nested within the picobirnavirus clade (Supplementary Figure 2).

**Figure 2:**
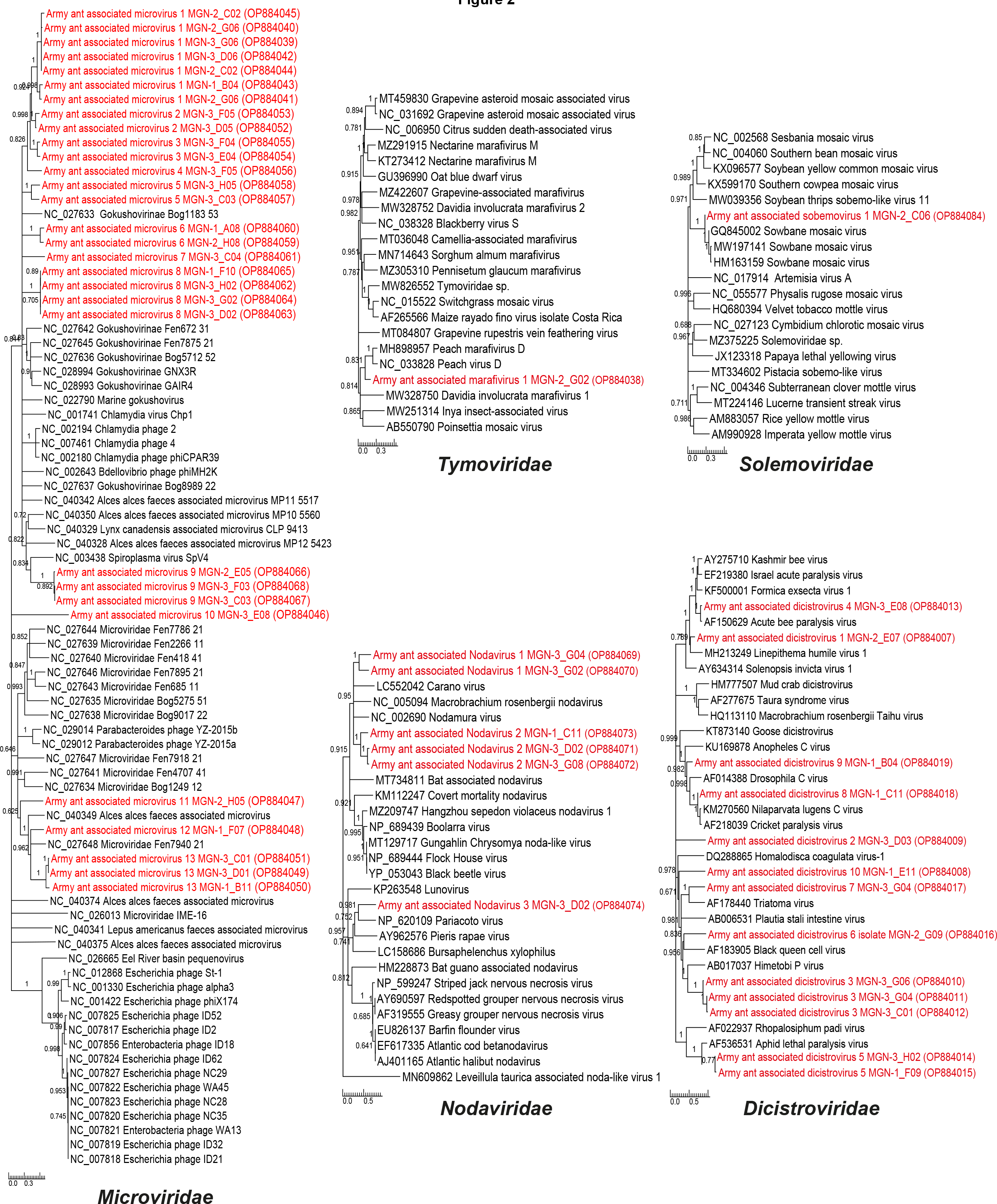
Phylogenies of the sequences of the viruses in the families *Microviridae*, *Solemoviridae*, *Tymoviridae, Nodaviridae* and *Dicistroviridae*. Sequences in red refer to army ant-associated contigs. Neighbor joining phylogenetic trees were generated using alignments of *Microviridae* major capsid protein sequences ranging from 306 aa to 761 aa, *Dicistroviridae* RNA-dependent RNA polymerase protein sequences (694-2775 aa), *Nodaviridae* RNA- dependent RNA polymerase protein sequences (656-1033 aa), *Solemoviridae* RNA-dependent RNA polymerase protein sequences (262 aa) and *Tymoviridae* polyprotein protein sequences (117 aa). One thousand bootstrap replicates were performed to quantify branch support. The scale bar depicts the number of amino acid substitutions per site.

#### Stramenopiles/alveolates/Rhizaria viruses

A total of 152 contigs ≥200 nt in length (mean size = 282 nt) were assigned to the crucivirus group (Supplementary Table 2): a growing group of viruses that appear to have originated through recombination between circular Rep-encoding ssDNA (CRESS-DNA) viruses and RNA tombusviruses that may infect members of the stramenopiles/alveolates/Rhizaria supergroup ^40^. Phylogenetic analyses of the predicted capsid protein sequences of three crucivirius-like contigs (for which we obtained CP sequences >440 aa) and representative sequences of cruciviruses showed that the ant-derived sequences clustered with two unclassified cruciviruses (Crucivirus 250 [Acc. Nbr: MT263579] and Crucivirus 268 [Acc. Nbr: MT263584]) isolated from water and sediments in New Zealand (Supplementary Figure 2).

#### Plant viruses

Overall, 101 contigs ≥200 nt in length were assigned to 22 genera in 14 plant virus families (Figure 1, Supplementary Table 2). The size of these contigs ranged from 203 nt to 745 nt (mean = 298 nt), suggesting that they may have originated from degraded plant virus nucleic acids. This degradation could likely be a consequence of the digestion process, either from plants directly consumed by the ants, or along the trophic chain that ended in their eventual presence within the sampled army ants. Alternatively, the low yield of reads/contigs assigned to plant viruses may have been due to the sequencing depth being insufficient for the detection of low abundance viruses within communities sometimes comprising ten or more virus species present at widely different titres. Plant virus contigs were recovered from 30/145 individual ants (20.7%), including 22/78 soldiers (28.2%) and 8/67 workers (11.9%), suggesting that, as with other viruses, plant viruses tended to accumulate more in soldiers. Contigs with detectable homology with plant virus families containing economically relevant crop pathogens (e.g., *Reoviridae*, *Tombusviridae*, *Geminiviridae*, *Solemoviridae*, or *Alphaflexiviridae* Figure 1) were identified in this study and in a recent study focusing on North American red fire ant *Solenopsis invicta* ^45^. This confirms that a broad diversity of plant viruses can potentially be detected within the viromes of top-end predators like army ants that feed on a wide range of herbivorous insects such as whiteflies, aphids, leafhoppers and thrips. Interestingly, possible translations of several contigs shared high amino acid identity (86-98%) with two well studied plant viruses: Peach virus D (PeVD; *Tymoviridae* family; *Marafivirus* genus) and Sowbane mosaic virus (SoMV; *Solemoviridae* family; *Sobemovirus* genus). While PeVD-like and SoMV-like contigs (≥ 200 nt) were respectively recovered from four and five ant colonies, smaller (100 - 200 nt) PeVD- like and SoMV-like contigs were, respectively identified from 11 and 9 colonies. These two plant viruses thus appear highly prevalent in the hunting areas of the sampled army ants. Mapping of Illumina and Nanopore reads and contigs assigned to PeVD and SoMV against reference genomes enabled the assembly of partial genome scaffolds respectively covering 39% (2576/6612 nt) and 79.4% (3163/3983 nt) of the PeVD and SoMV full length genomes. Phylogenetic analyses of the SoMV-like RdRp sequences and the PeVD-like polyprotein sequences confirmed that these army ant-associated viruses respectively clustered with known SoMV and PeVD isolates (Figure 2). Even though contigs assigned to 22 genera in 14 plant virus families were obtained from army ant samples, it is questionable whether these top-end predators are the best samplers for plant metavirome-focused studies. Using herbivorous insects such as caterpillars that directly feed on a wide variety of plants or using predators such as dragonflies^20^, damselflies, ^21^ or ladybug larva that prey on a wide variety of herbivorous insects, would probably be more efficient with respect to analysing the metaviromes of plants within an environment.

#### Invertebrate-infecting viruses

Besides recovering a diverse array of vertebrate-, plant- and prokaryote-associated virus sequences, we also identified a large number of highly diverse contigs with detectable homology to known invertebrate-infecting viruses. These included single-strand positive-sense RNA viruses of the *Nodaviridae* (82 contigs, mean size = 678 nt), *Dicistroviridae* (643 contigs, mean size = 496 nt) and *Iflaviridae* (1067 contigs, mean size = 549 nt) families (Supplementary Table 2 and Figures 2 and 3), branching with previously known viral sequences associated with or infecting invertebrates. We also found 1273 contigs sharing detectable homology with the order *Picornavirales* (Supplementary Table 2), including unclassified Picorna-like viruses (793 contigs, mean size = 609 nt) and viruses assigned to the *Sopolycivirus* and *Chipolycivirus* genera of the *Polycipiviridae* family (480 contigs, mean size = 481 nt; Figure 3), two genera of non-segmented, linear, positive-sense RNA viruses that have previously been found associated with ants (family *Formicidae*) ^52^. Finally, we recovered 103 contigs (mean size = 793 nt) that were assigned to the *Bidnaviridae* family: a virus family that is composed of bipartite ssDNA viruses infecting invertebrates. Interestingly, it has been suggested that bidnaviruses may have arisen by integration of an ancestral parvovirus genome into a large virus-derived DNA transposon from the polinton family ^53^. We here propose that the bidnavirus-like contigs recovered from army ants in our study are likely derived from diverse bipartite bidnaviruses, because these contigs are scattered all through the bidnavirus phylogenetic tree (Figure 3).

**Figure 3:**
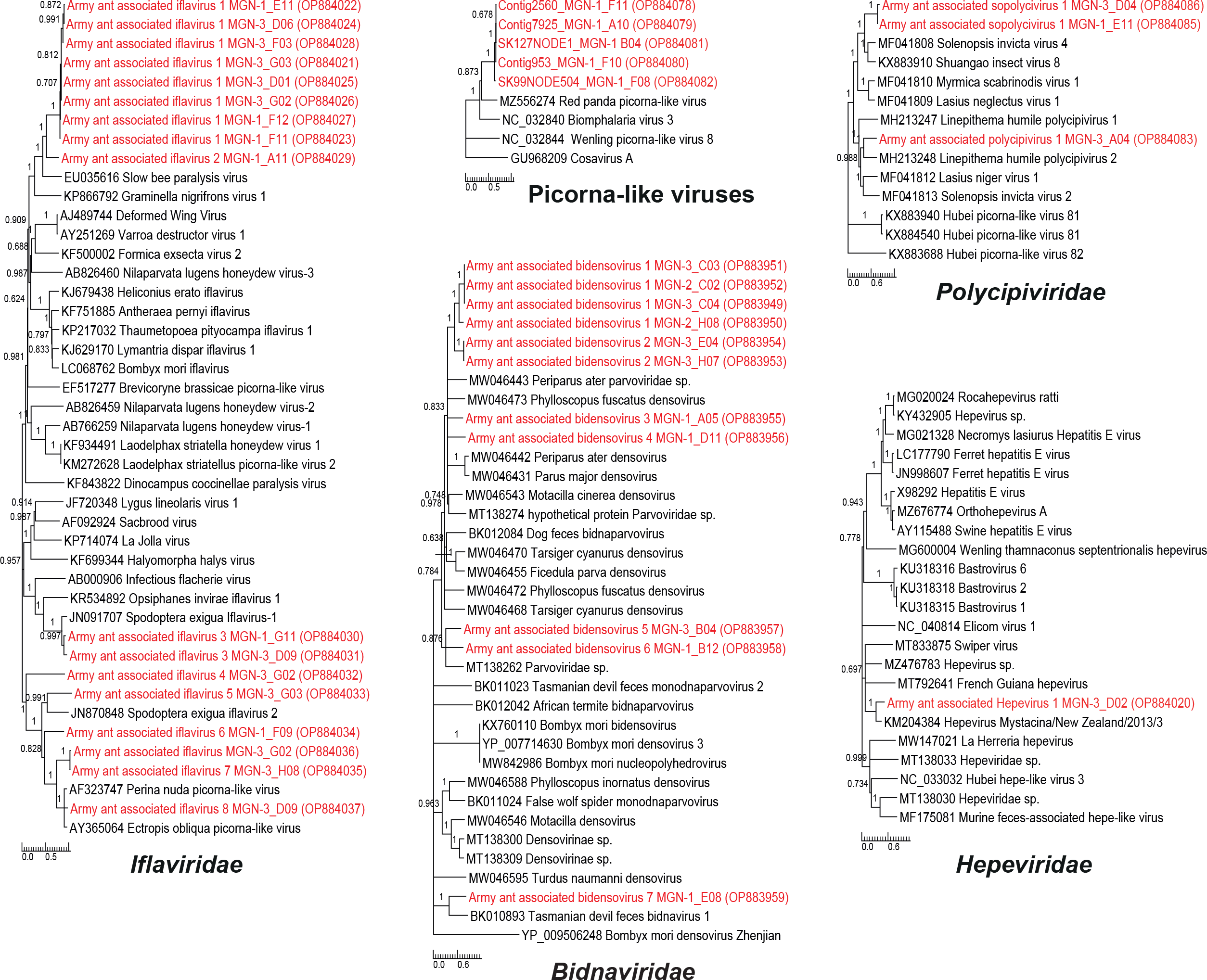
Phylogenies of the sequences of the viruses in the families *Iflaviridae*, *Bidnaviridae*, *Hepeviridae* and *Polycipiviridae*, as well as Picorna-like viruses. Sequences in red refer to army ant-associated contigs. Neighbor joining phylogenetic trees were generated using alignments of *Iflaviridae* polyprotein protein sequences ranging from 965 aa to 3229 aa, *Hepeviridae* polyprotein protein sequences (669 aa), Picorna-like viruses polyprotein protein sequences (2487-2858 aa), *Bidnaviridae* major capsid protein sequences (515-615 aa) and *Polycipiviridae* ORF5 protein sequences (1479-2331 aa) . One thousand bootstrap replicates were performed to quantify branch support. The scale bar depicts the number of amino acid substitutions per site.

#### Vertebrate-infecting viruses

Because *Dorylus* species have been reported to occasionally prey on vertebrates and scavenge on vertebrate carcasses, we logically investigated the presence of vertebrate viruses. Besides contigs assigned to cyclovirus and parvovirus taxonomic groups that are known to infect vertebrates (see below for a detailed analysis of these two taxa), we found 12 contigs that shared high nucleotide identity with the *Hepeviridae* family (mean size = 491 nt), including 4 contigs assigned to the *Orthohepevirus* genus (Supplementary Table 2). The family *Hepeviridae* includes five genera whose members infect salmonid fish, mammals and birds ^54^. Specifically, a 2017 nt long contig shared 72.73% identity (e-value = 1.10^-70^) with Hepevirus Mystacina/New Zealand/2013/3 (Acc. Nbr: KM204384), a hepevirus that was discovered associated with New Zealand lesser short-tailed bats (*Mystacina tuberculata*) ^55^. This result was further confirmed by phylogenetic analyses (Figure 3), and supports the hypothesis that the viromes of army ants can include viruses derived from the vertebrates upon which they prey and /or scavenge.

#### Foraging army ants are reservoirs of a large diversity of parvoviruses and cycloviruses

We focused on two viral families that have been extensively studied in recent years in the context of metagenomics studies: *Parvoviridae* and *Circoviridae*. Both are ssDNA virus families with member species sharing many biological, epidemiological and ecological characteristics. During the last decade, parvoviruses (in particular chapparvoviruses) and circoviruses (in particular cycloviruses) have been characterized from various fluid (e.g. blood) and excretion samples (e.g. feces) from vertebrates ^56–59^. Some of these studies have further demonstrated that some chapparvoviruses and cycloviruses can actually infect vertebrate hosts ^57–59^. However, due to the wide distribution of both viral families also in invertebrates, their natural host-ranges and the pathological characteristics of these viruses remain largely uncertain. For this reason, we were particularly interested in determining the phylogenetic relationships between parvovirus- and cyclovirus sequences found in army ants and those of close relatives that have been sampled previously from vertebrates and invertebrates.

#### Parvoviruses

The family *Parvoviridae* is composed of animal-infecting viruses which can collectively infect almost all major vertebrate clades and both proto- and deuterostome invertebrates ^60^. Parvovirus-related sequences (PRS) were by far the most abundant virus sequences amplified from the army ant samples and accounted for 17,419 contigs (77.8% of all virus contigs ≥200 nt in length, mean size = 469 nt).

Four hundred and three army ant-associated SF3 helicase domain sequences (a subdomain of the parvovirus NS1 protein) were scattered around the parvovirus SF3 phylogenetic tree and clustered with SF3 sequences from viruses in almost all parvovirus genera with the notable exception of those in genera of the *Parvovirinae* subfamily (Figure 4). This suggests that a range of animals infected by a broad diversity of parvoviruses was likely preyed or scavenged upon by army ant colonies. These animals were likely predominantly invertebrates including arthropods, molluscs, annelids, nematodes, and cnidarians (Figure 4). This phylogenetic analysis also revealed parvoviruses that might infect army ants. Specifically, some clades contained only army ant-derived SF3 sequences (see for instance the clade located between the two ambidensovirus groups, depicted with an ant and a question mark in Figure 4) and it is plausible that the parvoviruses from which these sequences were derived may have been directly infecting the army ants.

**Figure 4:**
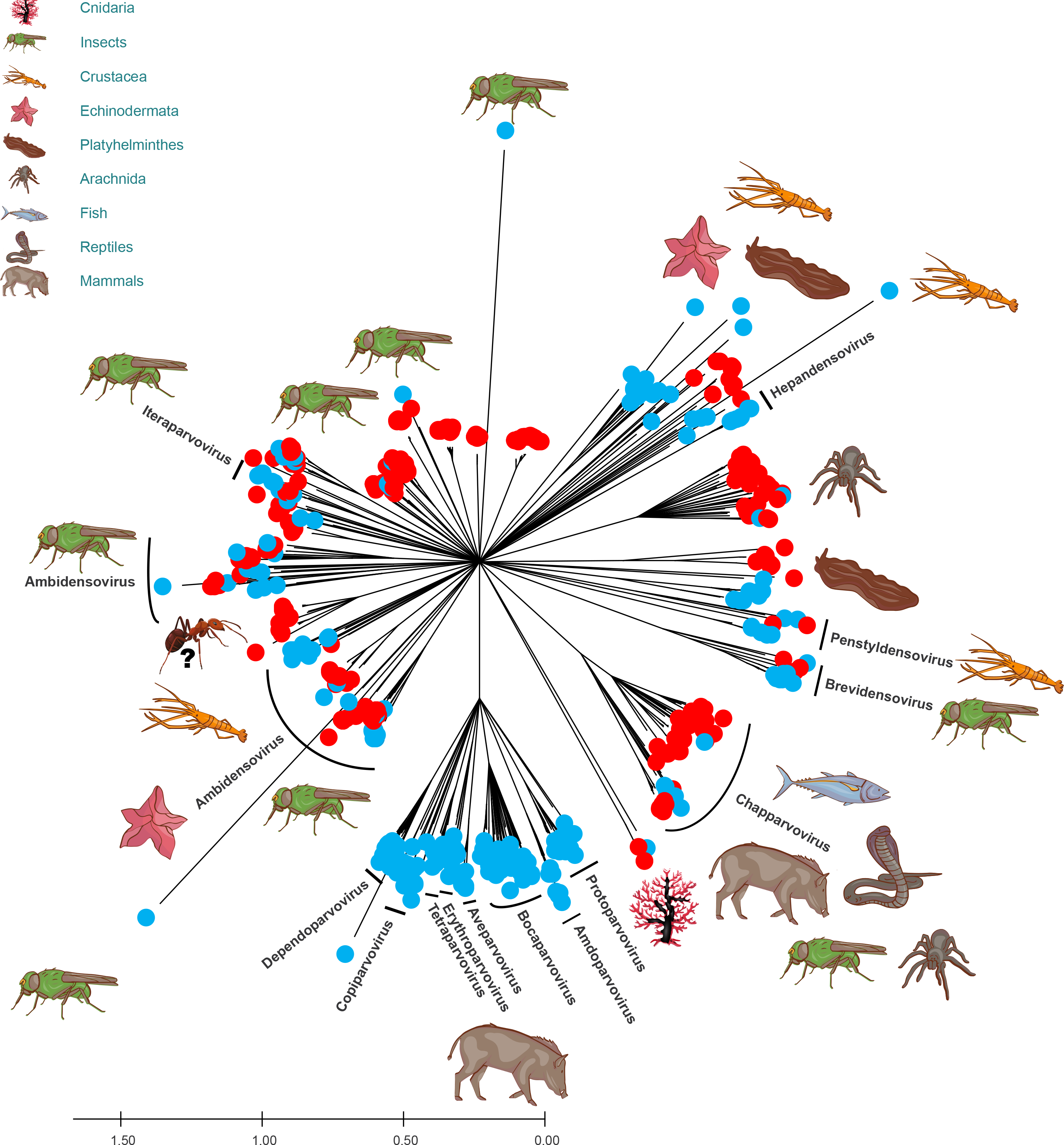
Phylogeny of the sequences of the viruses in the family *Parvoviridae*. The Neighbor joining phylogenetic tree was generated using alignments of SF3 protein sequences with 500 bootstrap replicates to quantify branch support. Sequences in red refer to army ant- associated contigs. Putative hosts of several parvoviruses are depicted at the extremity of the branches. Specifically, an ant and a question mark are depicted nearby the clade located between both ambidensovirus groups, because it is plausible that the parvoviruses from which these sequences were derived may have been directly infecting the army ants. Genera of the *Parvoviridae* family are also indicated. The scale bar depicts the number of amino acid substitutions per site.

We further focused on the genetic relationships of the NS1 proteins of chapparvovirus isolates because, while these parvoviruses have been primarily identified by metagenomic studies of animal feces, they have also been both isolated from the tissues of vertebrates (including reptiles, mammals, and birds ^56, 61^) and found as endogenous parvoviral elements (EPVs) within invertebrate genomes ^62^. Recently, this parvovirus group was split into two distinct sub-groups corresponding to two newly established genera: *Ichthamaparvovirus* and *Chaphamaparvovirus*^60^. While ichthamaparvoviruses are known to infect fishes and potentially also invertebrates, chaphamaparvoviruses have only so far been found in vertebrates ^62^.

Among the parvovirus-like contigs from army ants were several lineages of highly diverse chapparvoviruses (Figure 5) that likely reflect the wide range of prey and carrion upon which these ants feed. These apparently chapparvovirus-derived contigs clustered with invertebrate EPVs (*Mesobuthus martensii*, *Catajapyx aquilonaris* and *Nephila pilipes*) and sequences recovered from bird swabs and a human plasma sample (Figure 5). This suggests that the diversity of chapparvoviruses that infect army ants potentially resulted from the ingestion of infected invertebrates and that chapparvoviruses isolated from animal feces or human plasma may have been derived directly from either parasitic or ingested invertebrates

**Figure 5:**
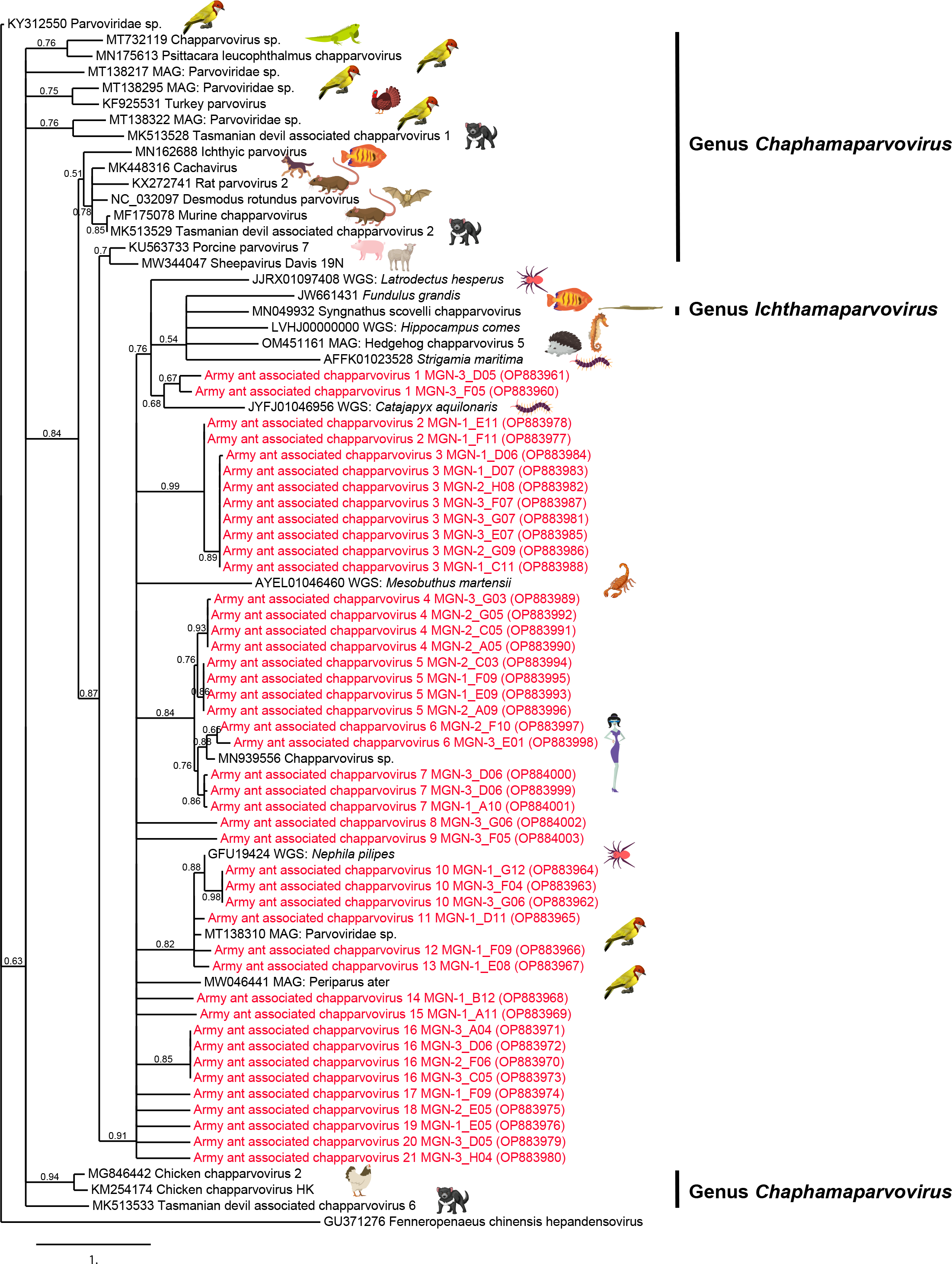
Phylogeny of the chapparvoviruses. The maximum likelihood tree was generated using alignments of NS1 protein sequences ranging from 214 aa to 823 aa and the WAG substitution model assuming an estimated proportion of invariant sites (of 0.075) and 4 gamma- distributed rate categories to account for rate heterogeneity across sites. The gamma shape parameter was estimated directly from the data (gamma=1.337). Support for internal branches was assessed using the aLRT test (SH-Like). Sequences in red refer to army ant-associated contigs. Putative hosts of several parvoviruses are depicted to the right of the branches. The two genera of the chapparvovirus group are also indicated. The scale bar depicts the number of amino acid substitutions per site.

#### Cycloviruses

Four hundred and seventy-two contigs ≥ 200 nt in length (mean size = 417 nt) with detectable homology to sequences of viruses in the family *Circoviridae* (that comprises two genera: *Circovirus* and *Cyclovirus*) clustered in 22 genetic groups (data not shown). Here, we were able to obtain potentially complete genomes of 45 isolates representing the 22 genetic groups from 17 army ant colonies. These genomes, with lengths ranging from 1,723 to 2,024 nt, contained two ORFs predicted to encode a replication-associated protein (Rep) and a capsid (CP) protein. A conserved nonanucleotide motif (TAGTATTAC) that is typical of the members of the *Circoviridae* family was identified on the *cp*-encoding strand of the 45 genomic sequences. Similarly, all genomes contained two intergenic regions, one located between the 5′ ends of the *rep* and *cp* ORFs, and another between the 3′ ends of the *cp* and *rep* ORFs: again genomic features common in the *Circoviridae* family. Finally, ten out of the 45 *Circoviridae* genome sequences have an apparent intron in the *rep* coding region that is possibly spliced to yield a functional Rep. A phylogenetic analysis showed that the 45 translated Rep sequences are all nested within the phylogenetic tree that included Rep sequences of representative isolates of the *Cyclovirus* genus (Figure 6). Thirty-eight out of the 45 army ant-associated cyclovirus Rep sequences clustered in four groups that respectively contained seven isolates (cluster I in Figure 6), eleven isolates (cluster II in Figure 6), ten isolates (cluster III in Figure 6) and ten isolates (cluster VI in Figure 6). Given that viruses within the *Circoviridae* family are classified into species based on genome-wide pairwise identities with an 80% species demarcation threshold ^63^, the 45 army ant-associated cycloviruses could reasonably be classified into nine new species (Army ant associated cyclovirus 1, 2, 3, 4, 5, 6, 8, 9 and 10) and two existing *Cyclovirus* species.

**Figure 6:**
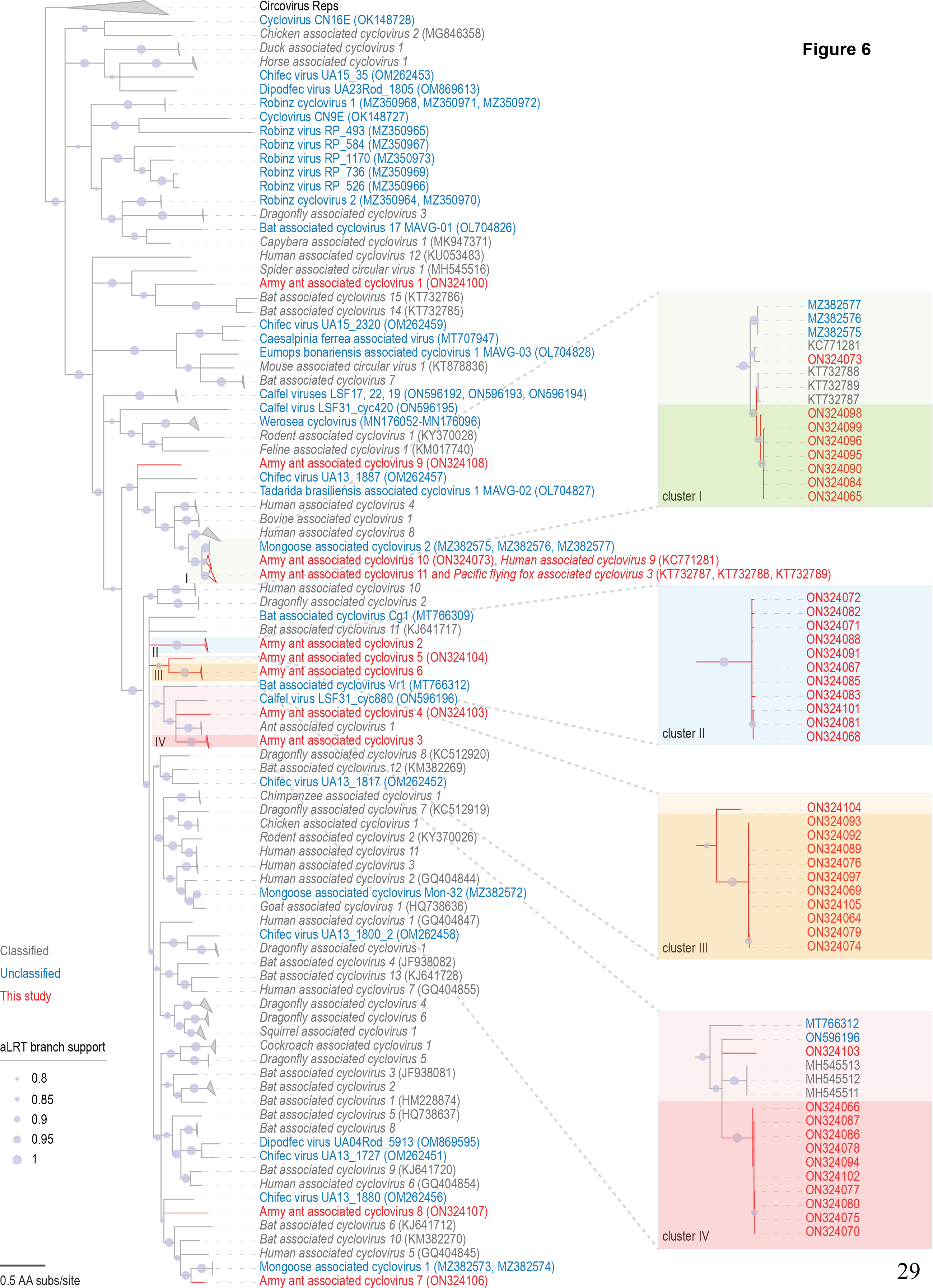
Phylogeny of the cycloviruses. The maximum likelihood tree was generated using alignments of Rep protein sequences and the LG+I+G substitution model. Support for internal branches was assessed using the aLRT test (SH-Like). Branches in red, grey and blue refer to army ant-associated, classified and unclassified cyclovirus Rep sequences. The scale bar depicts the number of amino acid substitutions per site. Branches with <0.8 aLRT support have been collapsed with TreeGraph 2 ^68^. Zoomed-in views of four regions of the phylogenetic tree are depicted on the right side of the figure to better localize isolates of Cluster I (dark green area), Cluster II (light blue area), Cluster III (dark orange area) and Cluster IV (dark red area).

Specifically, the seven isolates of cluster I (Army ant associated cyclovirus 11) shared 90.8- 91.8% identity with Pacific flying fox associated cyclovirus 3 (Acc. Nbr. KT732787, KT732788 and KT732789) and Army ant associated cyclovirus 7 (Acc. Nbr ON324106) shared 80.6% with Bat cyclovirus isolate CyVLysokaP4_CMR_2014 (Acc. Nbr MG693174) which was isolated in Cameroon from fecal samples of *Eidolon helvum*, a fruit-eating bat ^64^. Interestingly, a Rep sequence recovered from one of the army ant samples (Army ant associated cyclovirus 10, Acc. Nbr. ON324073) shared 262/278 amino acids (94% identity) with a human cyclovirus (isolate VS5700009, Acc. Nbr. YP_008130363) isolated from a patient with an unexplained paraplegia from Malawi ^59^. The CP sequence predicted from the same assembled genome-length contig shared 162/185 aa (88% identity) with an unclassified cyclovirus (isolate ZM36a, Acc. Nbr. YP_009104365) detected within the intestinal contents of a shrew in Zambia ^65^. Cyclovirus ZM36a was also closely related to cycloviruses initially identified from human patients with central nervous system manifestations ^66^. These results stress the need to better monitor and understand the circulation of cycloviruses between invertebrates, humans and rodents, and the potential of army ants as natural samplers: perhaps even of viruses directly related to human diseases.

## Conclusion

This study suggests that predators and scavengers such as army ants can be used to sample broad swathes of environmental viromes including viruses infecting plants, invertebrates and vertebrates. Although not completely unbiased (insects will still have feeding preferences that preclude the sampling of all viruses in an ecosystem), using army ants to sample tropical forest viromes will likely yield sequence data from a more diverse array of viruses than if plants or animals with less diverse diets than army ants were sampled. Army ants will be a particularly good tool for sampling invertebrate viruses in such environments given that they carry what appears to be an extraordinary diversity and abundance of sequences related to known invertebrate-infecting viruses. Although how thoroughly army ants sample the complete invertebrate-associated viromes in the areas surrounding their temporary nests is still unclear, it is undisputable that these top-end predators are probably sampling a non-negligible fraction thereof.

Longitudinal metagenomic analyses of army ant-associated viral nucleic acids in agro- ecological interfaces such as tropical forest areas that bound managed farmlands or human settlements could be a highly convenient means of gaining insights into the ecosystem-scale impacts of natural or human-mediated environmental changes on virus population compositions and structure. Conversely, monitoring of changes in the relative diversity and prevalence of different viral lineages within the invertebrate-associated virome (and even within plant- and vertebrate-associated viromes) over time could provide a sensitive “leading” indicator of changes in tropical forest ecosystem stability that could foreshadow more obvious changes in these ecosystems due to climate change and other human-mediated disturbances ^67^.

## Acknowledgements

This work has been realized with the support of MESO@LR-Platform at the University of Montpellier.

## Funding

The study was funded by the Word Organization for Animal Health (EBO-SURSY: FOOD/2016/379-660)

## Conflict of interest disclosure

The authors declare that they comply with the PCI rule of having no financial conflicts of interest in relation to the content of the article.

## Author contributions

S.B., E.L. and P.R. designed the experiment; M.F., P.B., T.N.M., L.B.K., L.H.L., L.B., I.M.M, G.D.M, F.R.N, J.S.K, M.Y, S.B and E.L collected the army ants; M.F., B.R., L.C., E.F., S.K. and J.M.C. performed the molecular biology experiments; M.F., B.R., D.F., F.M., M.O., A.V. and P.R. analyzed the data; B.R., A.V. and P.R. produced the figures; D.P.M. and P.R. wrote the first draft, all authors contributed editorially to the manuscript, and author order was determined on the basis of these duties.

## Data, scripts, code, and supplementary information availability

The cleaned sequence read data from the Illumina HiSeq runs and the raw sequence read data from the Flongle Nanopore runs are available from the NCBI Sequence Read Archive, under the Bioproject accession number PRJNA900057 (Biosample accession numbers SAMN31677877, SAMN31677878, SAMN31677879, SAMN31726898, SAMN31726899, SAMN31726900 and SAMN31726901) and the following URLs:(https://www.ncbi.nlm.nih.gov/sra/?term=SRR22334884; https://www.ncbi.nlm.nih.gov/sra/?term=SRR22334885; https://www.ncbi.nlm.nih.gov/sra/?term=SRR22334886; https://www.ncbi.nlm.nih.gov/sra/?term=SRR22334634; https://www.ncbi.nlm.nih.gov/sra/?term=SRR22334635; https://www.ncbi.nlm.nih.gov/sra/?term=SRR22334636; https://www.ncbi.nlm.nih.gov/sra/?term=SRR22334637). In addition, the sequence reads per sample are available from the corresponding author. The 138 army ant associated contig sequences used in the phylogenetic analyses were deposited in GenBank under accession numbers OP883949- OP884086. The 45 army ant associated cyclovirus genome sequences were deposited in GenBank under accession numbers ON324064-ON324108. Script and codes were deposited in Zenodo with the following DOI: https://doi.org/10.5281/zenodo.7636216

## SUPPLEMENTARY MATERIAL

### Supplementary figure legends

**Supplementary Figure 1:**
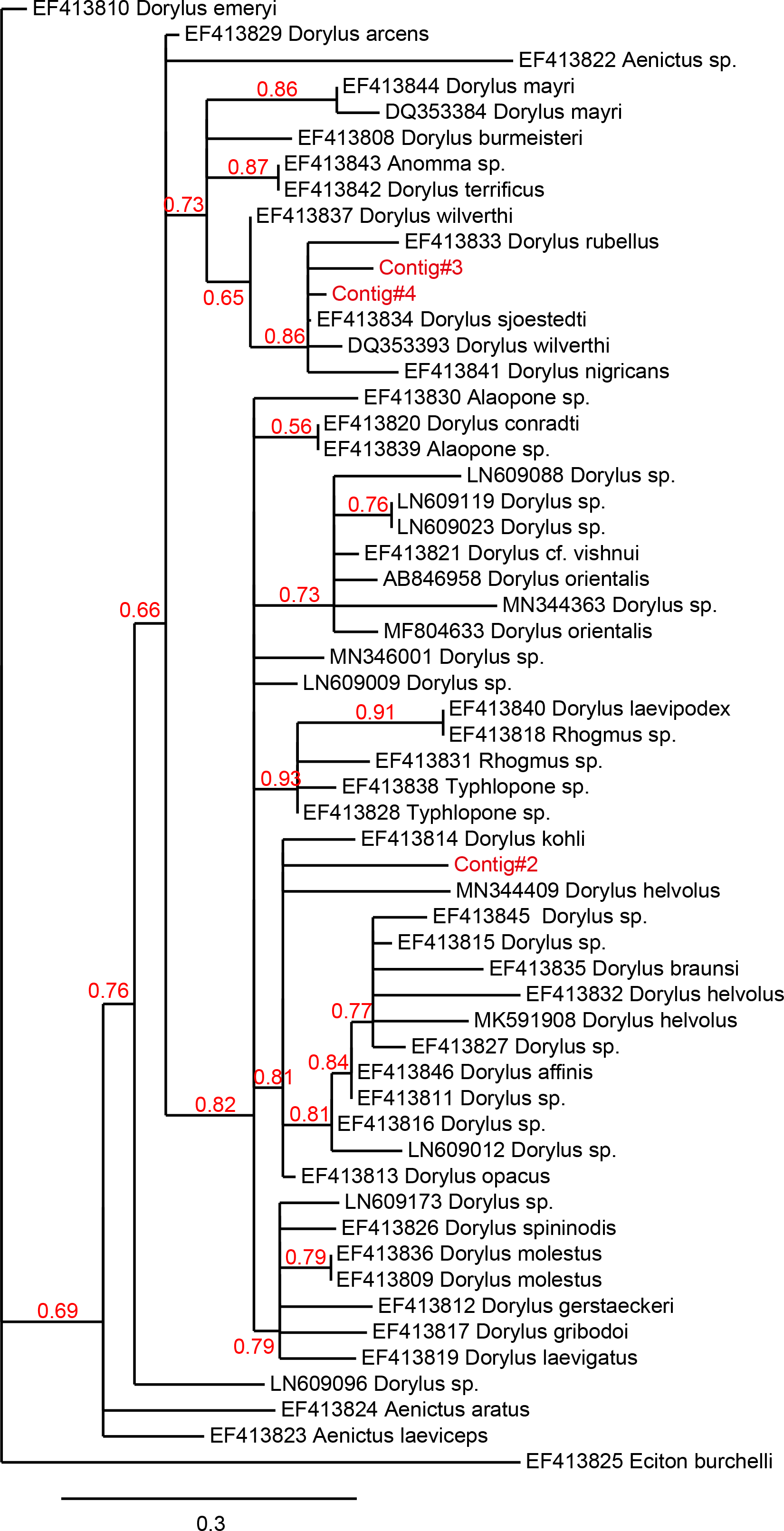
Phylogeny of the Illumina reads assigned to *Dorylus* sp. cytochrome oxidase I gene using BLASTx searches. A phylogenetic tree was constructed using the maximum likelihood method using the HKY85 substitution model assuming an estimated proportion of invariant sites of 0.542 and 4 gamma-distributed rate categories to account for rate heterogeneity across sites. The gamma shape parameter was estimated directly from the data (gamma=1.219). Support for internal branches was assessed using the aLRT test (SH- Like).

**Supplementary Figure 2:**
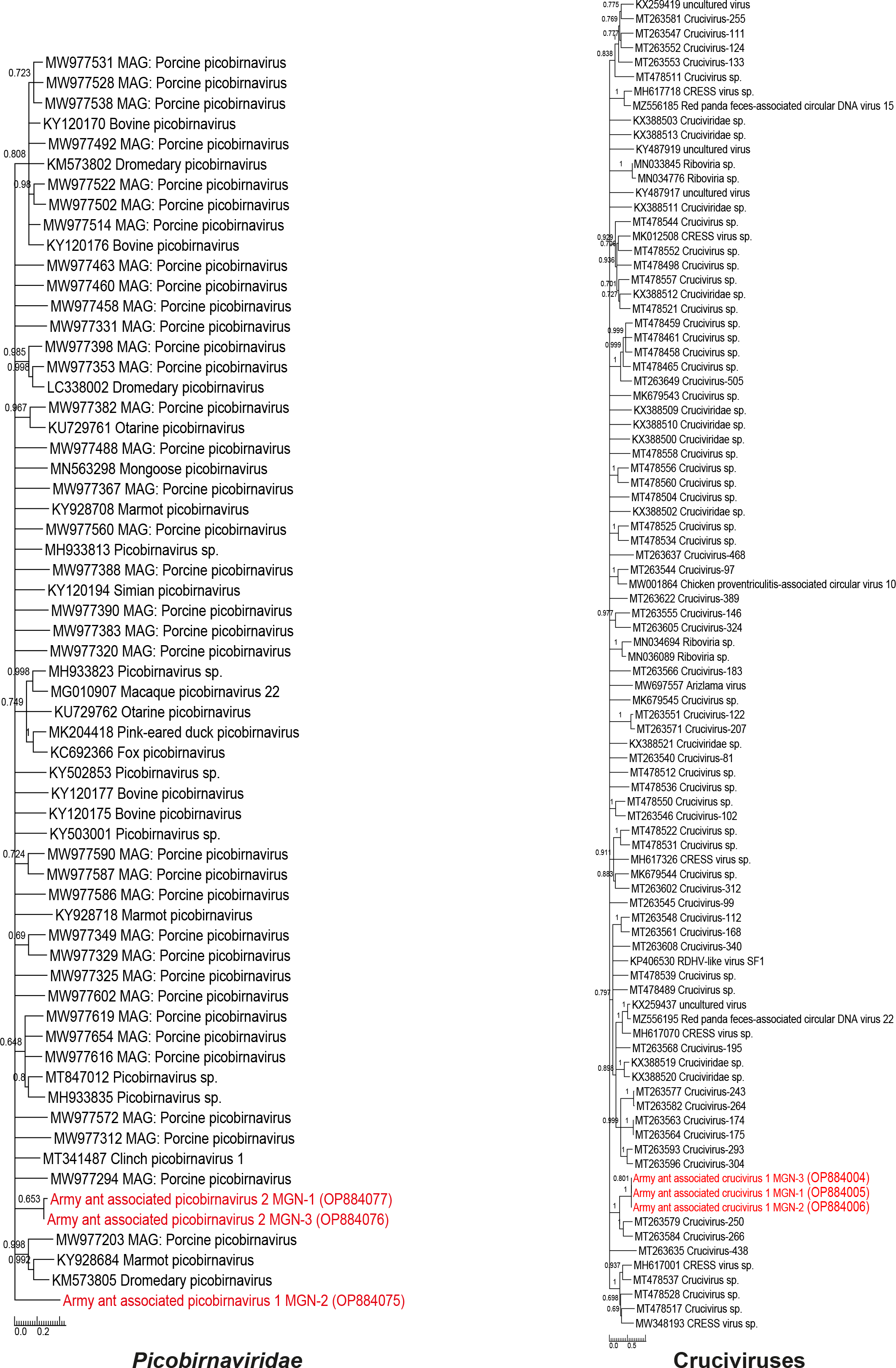
Phylogenies of the sequences of the viruses in the *Picobirnaviridae* family, as well as cruciviruses. Sequences in red refer to army ant associated contigs. Neighbor joining phylogenetic trees were generated using alignments of capsid protein sequences of cruciviruses (444 aa) and *Picobirnaviridae* RNA-dependent RNA polymerase protein sequences (137-201 aa). One thousand bootstrap replicates were performed to quantify branch support. The scale bar depicts the number of amino acid substitutions per site.

### Supplementary table legends

**Supplementary Table 1:** Additional characteristics of army ant samples. “nd” means not determined.

| Ant colony number | Sample number | Sequencing number | Sample taxonomy | Number of processed ants | Genbank Accession Number |
| --- | --- | --- | --- | --- | --- |
| FM1 | 185 | MGN-3_B02 | *D. sjoestedti/wilverthi* clade | 1 worker, 1 soldier | SRX18307458 |
| FM1 | 186 | MGN-3_C02 | nd | 1 worker, 1 soldier | SRX18307707 |
| FM1 | 183 | MGN-3_G03 | nd | 1 worker, 1 soldier | SRX18307707 |
| FM1 | 184 | MGN-3_H03 | nd | 1 worker, 1 soldier | SRX18307707 |
| FM2 | 165 | MGN-3_A04 | nd | 1 worker, 1 soldier | SRX18307707 |
| FM2 | 166 | MGN-3_B04 | *D. sjoestedti/wilverthi* clade | 1 worker, 1 soldier | SRX18307707 and SRX18307457 |
| FM2 | 163 | MGN-3_G05 | *D. sjoestedti/wilverthi* clade | 1 worker, 1 soldier | SRX18307707 |
| FM2 | 164 | MGN-3_H05 | nd | 1 worker, 1 soldier | SRX18307707 |
| FM3 | 188 | MGN-3_D02 | nd | 1 worker, 1 soldier | SRX18307707 |
| FM3 | 189 | MGN-3_E02 | nd | 1 worker, 1 soldier | SRX18307707 |
| FM3 | 190 | MGN-3_F02 | nd | 1 worker, 1 soldier | SRX18307707 |
| FM3 | 191 | MGN-3_G02 | nd | 1 worker, 1 soldier | SRX18307707 |
| FM4 | 194 | MGN-3_B01 | nd | 1 worker, 1 soldier | SRX18307707 |
| FM4 | 195 | MGN-3_C01 | nd | 1 worker, 1 soldier | SRX18307707 |
| FM4 | 196 | MGN-3_D01 | nd | 1 worker, 1 soldier | SRX18307707 |
| FM4 | 193 | MGN-3_H02 | *D. sjoestedti/wilverthi* clade | 1 worker, 1 soldier | SRX18307707 |
| FM5 | 175 | MGN-3_A03 | *D. sjoestedti/wilverthi* clade | 1 worker, 1 soldier | SRX18307707 |
| FM5 | 176 | MGN-3_B03 | nd | 1 worker, 1 soldier | SRX18307707 |
| FM5 | 173 | MGN-3_G04 | *D. sjoestedti/wilverthi* clade | 1 worker, 1 soldier | SRX18307707 |
| FM5 | 174 | MGN-3_H04 | *D. sjoestedti/wilverthi* clade | 1 worker, 1 soldier | SRX18307707 |
| FM6 | 178 | MGN-3_C03 | nd | 1 worker, 1 soldier | SRX18307707 |
| FM6 | 179 | MGN-3_D03 | nd | 1 worker, 1 soldier | SRX18307707 |
| FM6 | 180 | MGN-3_E03 | nd | 1 worker, 1 soldier | SRX18307707 |
| FM6 | 181 | MGN-3_F03 | nd | 1 worker, 1 soldier | SRX18307707 |
| FM7 | 168 | MGN-3_C04 | *D. sjoestedti/wilverthi* clade | 1 worker, 1 soldier | SRX18307707 |
| FM7 | 169 | MGN-3_D04 | nd | 1 worker, 1 soldier | SRX18307707 |
| FM7 | 170 | MGN-3_E04 | *D. sjoestedti/wilverthi* clade | 1 worker, 1 soldier | SRX18307707 |
| FM7 | 171 | MGN-3_F04 | nd | 1 worker, 1 soldier | SRX18307707 |
| FM8 | 198 | MGN-3_E01 | nd | 1 worker, 1 soldier | SRX18307707 |
| FM8 | 199 | MGN-3_F01 | nd | 1 worker, 1 soldier | SRX18307707 |
| FM8 | 200 | MGN-3_G01 | nd | 1 worker, 1 soldier | SRX18307707 |
| FM8 | 201 | MGN-3_H01 | nd | 1 worker, 1 soldier | SRX18307707 |
| P10 | P10-reste | MGN-1_A05 | nd | 4 workers, 4 soldiers | SRX18307705 |
| P10 | P10-1.1 | MGN-1_A06 | nd | 1 worker | SRX18307705 |
| P10 | P10-1.2 | MGN-1_B06 | nd | 1 soldier | SRX18307705 |
| P10 | P10-2.1 | MGN-1_C06 | nd | 1 worker | SRX18307705 |
| P10 | P10-2.2 | MGN-1_D06 | nd | 1 soldier | SRX18307705 |
| P10 | P10-3.1 | MGN-1_E06 | nd | 1 worker | SRX18307705 |
| P10 | P10-3.2 | MGN-1_F06 | nd | 1 soldier | SRX18307705 |
| P10 | P10-4.1 | MGN-1_G06 | nd | 1 worker | SRX18307705 |
| P10 | P10-4.2 | MGN-1_H06 | nd | 1 soldier | SRX18307705 |
| P11 | P11-4.2 | MGN-1_A04 | nd | 1 soldier | SRX18307705 |
| P11 | P11-reste | MGN-1_B04 | nd | 4 workers, 4 soldiers | SRX18307705 |
| P11 | P11-1.1 | MGN-1_B05 | nd | 1 worker | SRX18307705 |
| P11 | P11-1.2 | MGN-1_C05 | nd | 1 soldier | SRX18307705 |
| P11 | P11-2.1 | MGN-1_D05 | nd | 1 worker | SRX18307705 |
| P11 | P11-2.2 | MGN-1_E05 | nd | 1 soldier | SRX18307705 |
| P11 | P11-3.1 | MGN-1_F05 | nd | 1 worker | SRX18307705 |
| P11 | P11-3.2 | MGN-1_G05 | nd | 1 soldier | SRX18307705 |
| P11 | P11-4.1 | MGN-1_H05 | nd | 1 worker | SRX18307705 |
| P12 | P12-1.1 | MGN-1_C04 | nd | 1 worker | SRX18307705 |
| P12 | P12-1.2 | MGN-1_D04 | nd | 1 soldier | SRX18307705 |
| P12 | P12-2.1 | MGN-1_E04 | nd | 1 worker | SRX18307705 |
| P12 | P12-2.2 | MGN-1_F04 | *D. sjoestedti/wilverthi* clade | 1 soldier | SRX18307705 |
| P12 | P12-3.1 | MGN-2_A07 | nd | 1 worker | SRX18307706 |
| P12 | P12-3.2 | MGN-2_B07 | nd | 1 soldier | SRX18307706 |
| P12 | P12-4.1 | MGN-2_C07 | nd | 1 worker | SRX18307706 |
| P12 | P12-4.2 | MGN-2_D07 | nd | 1 soldier | SRX18307706 |
| P12 | P12-reste | MGN-2_E07 | nd | 4 workers, 4 soldiers | SRX18307706 |
| P13 | P13-2.2 | MGN-2_A06 | nd | 1 soldier | SRX18307706 |
| P13 | P13-3.1 | MGN-2_B06 | nd | 1 worker | SRX18307706 |
| P13 | P13-3.2 | MGN-2_C06 | nd | 1 soldier | SRX18307706 |
| P13 | P13-4.1 | MGN-2_E06 | nd | 1 worker | SRX18307706 |
| P13 | P13-4.2 | MGN-2_F06 | nd | 1 soldier | SRX18307706 |
| P13 | P13-1.1 | MGN-2_F07 | nd | 1 worker | SRX18307706 |
| P13 | P13-reste | MGN-2_G06 | nd | 4 workers, 4 soldiers | SRX18307706 |
| P13 | P13-1.2 | MGN-2_G07 | nd | 1 soldier | SRX18307706 |
| P13 | P13-2.1 | MGN-2_H07 | nd | 1 worker | SRX18307706 |
| P14 | P14-1.2 | MGN-2_A05 | nd | 1 soldier | SRX18307706 |
| P14 | P14-2.1 | MGN-2_B05 | nd | 1 worker | SRX18307706 |
| P14 | P14-2.2 | MGN-2_C05 | *D. sjoestedti/wilverthi* clade | 1 soldier | SRX18307706 |
| P14 | P14-3.1 | MGN-2_D05 | nd | 1 worker | SRX18307706 |
| P14 | P14-3.2 | MGN-2_E05 | nd | 1 soldier | SRX18307706 |
| P14 | P14-4.1 | MGN-2_F05 | nd | 1 worker | SRX18307706 |
| P14 | P14-4.2 | MGN-2_G05 | nd | 1 soldier | SRX18307706 |
| P14 | P14-reste | MGN-2_H05 | nd | 4 workers, 4 soldiers | SRX18307706 |
| P14 | P14-1.1 | MGN-2_H06 | nd | 1 worker | SRX18307706 |
| P15 | P15-reste | MGN-2_A03 | *D. sjoestedti/wilverthi* clade | 4 workers, 4 soldiers | SRX18307706 |
| P15 | P15-1.1 | MGN-2_A04 | nd | 1 worker | SRX18307706 |
| P15 | P15-1.2 | MGN-2_B04 | nd | 1 soldier | SRX18307706 |
| P15 | P15-2.1 | MGN-2_C04 | nd | 1 worker | SRX18307706 |
| P15 | P15-2.2 | MGN-2_D04 | nd | 1 soldier | SRX18307706 |
| P15 | P15-3.1 | MGN-2_E04 | nd | 1 worker | SRX18307706 |
| P15 | P15-3.2 | MGN-2_F04 | nd | 1 soldier | SRX18307706 |
| P15 | P15-4.1 | MGN-2_G04 | nd | 1 worker | SRX18307706 |
| P15 | P15-4.2 | MGN-2_H04 | nd | 1 soldier | SRX18307706 |
| P16 | P16-4.1 | MGN-2_A02 | nd | 1 worker | SRX18307706 |
| P16 | P16-4.2 | MGN-2_B02 | nd | 1 soldier | SRX18307706 |
| P16 | P16-1.1 | MGN-2_B03 | nd | 1 worker | SRX18307706 |
| P16 | P16-reste | MGN-2_C02 | *D. sjoestedti/wilverthi* clade | 4 workers, 4 soldiers | SRX18307706 |
| P16 | P16-1.2 | MGN-2_C03 | nd | 1 soldier | SRX18307706 |
| P16 | P16-2.1 | MGN-2_E03 | nd | 1 worker | SRX18307706 |
| P16 | P16-2.2 | MGN-2_F03 | nd | 1 soldier | SRX18307706 |
| P16 | P16-3.1 | MGN-2_G03 | nd | 1 worker | SRX18307706 |
| P16 | P16-3.2 | MGN-2_H03 | nd | 1 soldier | SRX18307706 |
| P17 | P17-1.1 | MGN-2_D02 | nd | 1 worker | SRX18307706 |
| P17 | P17-1.2 | MGN-2_E02 | nd | 1 soldier | SRX18307706 |
| P17 | P17-2.1 | MGN-2_F02 | *D. sjoestedti/wilverthi* clade | 1 worker | SRX18307706 |
| P17 | P17-2.2 | MGN-2_G02 | *D. sjoestedti/wilverthi* clade | 1 soldier | SRX18307706 and SRX18307455 |
| P17 | P17-3.1 | MGN-2_H02 | nd | 1 worker | SRX18307706 |
| P17 | P17-3.2 | MGN-3_A09 | nd | 1 soldier | SRX18307707 |
| P17 | P17-4.1 | MGN-3_B09 | nd | 1 worker | SRX18307707 |
| P17 | P17-4.2 | MGN-3_C09 | nd | 1 soldier | SRX18307707 |
| P17 | P17-reste | MGN-3_D09 | nd | 5 workers, 5 soldiers | SRX18307707 |
| P18 | P18-3.1 | MGN-3_A08 | nd | 1 worker | SRX18307707 |
| P18 | P18-3.2 | MGN-3_B08 | nd | 1 soldier | SRX18307707 |
| P18 | P18-4.1 | MGN-3_C08 | nd | 1 worker | SRX18307707 |
| P18 | P18-4.2 | MGN-3_D08 | nd | 1 soldier | SRX18307707 |
| P18 | P18-reste | MGN-3_E08 | *D. sjoestedti/wilverthi* clade | 4 workers, 4 soldiers | SRX18307707 |
| P18 | P18-1.1 | MGN-3_E09 | nd | 1 worker | SRX18307707 |
| P18 | P18-1.2 | MGN-3_F09 | nd | 1 soldier | SRX18307707 |
| P18 | P18-2.1 | MGN-3_G09 | nd | 1 worker | SRX18307707 |
| P18 | P18-2.2 | MGN-3_H09 | nd | 1 soldier | SRX18307707 |
| P19 | P19-2.1 | MGN-3_A07 | nd | 1 soldier | SRX18307707 |
| P19 | P19-2.2 | MGN-3_B07 | nd | 1 soldier | SRX18307707 |
| P19 | P19-3.1 | MGN-3_C07 | nd | 1 soldier | SRX18307707 |
| P19 | P19-3.2 | MGN-3_D07 | nd | 1 soldier | SRX18307707 |
| P19 | P19-4.1 | MGN-3_E07 | nd | 1 soldier | SRX18307707 |
| P19 | P19-4.2 | MGN-3_F07 | nd | 1 soldier | SRX18307707 |
| P19 | P19-reste | MGN-3_G07 | nd | 1 worker, 7 soldiers | SRX18307707 |
| P19 | P19-1.1 | MGN-3_G08 | nd | 1 soldier | SRX18307707 |
| P19 | P19-1.2 | MGN-3_H08 | nd | 1 soldier | SRX18307707 |
| P1A | P1A-reste | MGN-1_A11 | nd | 4 workers, 4 soldiers | SRX18307705 |
| P1A | P1A-1.1 | MGN-1_A12 | nd | 1 worker | SRX18307705 |
| P1A | P1A-1.2 | MGN-1_B12 | nd | 1 soldier | SRX18307705 |
| P1A | P1A-2.1 | MGN-1_C12 | nd | 1 worker | SRX18307705 |
| P1A | P1A-2.2 | MGN-1_D12 | nd | 1 soldier | SRX18307705 |
| P1A | P1A-3.1 | MGN-1_E12 | nd | 1 worker | SRX18307705 |
| P1A | P1A-3.2 | MGN-1_F12 | nd | 1 soldier | SRX18307705 |
| P1A | P1A-4.1 | MGN-1_G12 | nd | 1 soldier | SRX18307705 |
| P1A | P1A-4.2 | MGN-1_H12 | nd | 1 worker | SRX18307705 |
| P21/23 | P21/23-1.2 | MGN-3_A06 | nd | 1 soldier | SRX18307707 |
| P21/23 | P21/23-2.1 | MGN-3_B06 | nd | 1 worker | SRX18307707 |
| P21/23 | P21/23-2.2 | MGN-3_C06 | *D. sjoestedti/wilverthi* clade | 1 soldier | SRX18307707 |
| P21/23 | P21/23-reste | MGN-3_D06 | *D. sjoestedti/wilverthi* clade | 5 workers, 5 soldiers | SRX18307707 |
| P21/23 | P21/23-1.1 | MGN-3_H07 | nd | 1 worker | SRX18307707 |
| P24 | P24-larves | MGN-3_A05 | nd | several larvae | SRX18307707 |
| P24 | P24-3.1 | MGN-3_B05 | nd | 1 worker | SRX18307707 |
| P24 | P24-3.2 | MGN-3_C05 | nd | 1 soldier | SRX18307707 |
| P24 | P24-4.1 | MGN-3_D05 | nd | 1 soldier | SRX18307707 |
| P24 | P24-4.2 | MGN-3_E05 | nd | 1 soldier | SRX18307707 |
| P24 | P24-1.1 | MGN-3_E06 | nd | 1 worker | SRX18307707 |
| P24 | P24-reste | MGN-3_F05 | nd | 4 workers, 4 soldiers | SRX18307707 |
| P24 | P24-1.2 | MGN-3_F06 | nd | 1 soldier | SRX18307707 |
| P24 | P24-2.1 | MGN-3_G06 | nd | 1 soldier | SRX18307707 |
| P24 | P24-2.2 | MGN-3_H06 | nd | 1 worker | SRX18307707 |
| P2A | P2A-1 | MGN-1_B11 | nd | 1 worker, 1 soldier | SRX18307705 |
| P2A | P2A-2 | MGN-1_C11 | *D. sjoestedti/wilverthi* clade | 1 worker, 1 soldier | SRX18307705 |
| P2A | P2A-3 | MGN-1_D11 | nd | 1 worker, 1 soldier | SRX18307705 |
| P2A | P2A-4 | MGN-1_E11 | nd | 1 worker, 1 soldier | SRX18307705 |
| P2A | P2A-reste | MGN-1_F11 | nd | 13 workers | SRX18307705 |
| P3A | P3A-1 | MGN-1_A10 | nd | 1 worker, 2 soldier | SRX18307705 |
| P3A | P3A-3.1 | MGN-1_B10 | nd | 2 workers | SRX18307705 |
| P3A | P3A-3.2 | MGN-1_C10 | nd | 1 soldier | SRX18307705 |
| P3A | P3A-4.1 | MGN-1_D10 | nd | 1 worker | SRX18307705 |
| P3A | P3A-4.2 | MGN-1_E10 | nd | 1 soldier | SRX18307705 |
| P3A | P3A-reste | MGN-1_F10 | nd | 4 workers, 4 soldiers | SRX18307705 |
| P3A | P3A-2.1 | MGN-1_G11 | nd | 1 worker | SRX18307705 |
| P3A | P3A-2.2 | MGN-1_H11 | *D. sjoestedti/wilverthi* clade | 1 soldier | SRX18307705 |
| P4A | P4A-2.1 | MGN-1_A09 | nd | 1 worker | SRX18307705 |
| P4A | P4A-2.2 | MGN-1_B09 | nd | 1 soldier | SRX18307705 |
| P4A | P4A-3.1 | MGN-1_C09 | nd | 1 worker | SRX18307705 |
| P4A | P4A-3.2 | MGN-1_D09 | nd | 1 soldier | SRX18307705 |
| P4A | P4A-4 | MGN-1_E09 | nd | 1 worker, 1 soldier | SRX18307705 |
| P4A | P4A-reste | MGN-1_F09 | *D. sjoestedti/wilverthi* clade | 4 workers, 4 soldiers | SRX18307705 |
| P4A | P4A-1.1 | MGN-1_G10 | nd | 1 worker | SRX18307705 |
| P4A | P4A-1.2 | MGN-1_H10 | *D. sjoestedti/wilverthi* clade | 1 soldier | SRX18307705 |
| P5 | P5-2 | MGN-1_A08 | nd | 1 worker, 1 soldier | SRX18307705 |
| P5 | P5-3.1 | MGN-1_B08 | nd | 1 worker | SRX18307705 |
| P5 | P5-3.2 | MGN-1_C08 | nd | 1 soldier | SRX18307705 |
| P5 | P5-4.1 | MGN-1_D08 | nd | 1 worker | SRX18307705 |
| P5 | P5-4.2 | MGN-1_E08 | *D. sjoestedti/wilverthi* clade | 1 soldier | SRX18307705 |
| P5 | P5-reste | MGN-1_F08 | nd | 4 workers, 4 soldiers | SRX18307705 |
| P5 | P5-1.1 | MGN-1_G09 | nd | 1 worker | SRX18307705 |
| P5 | P5-1.2 | MGN-1_H09 | nd | 1 soldier | SRX18307705 |
| P6 | P6-2.1 | MGN-1_A07 | nd | 1 worker | SRX18307705 |
| P6 | P6-2.2 | MGN-1_B07 | nd | 1 soldier | SRX18307705 |
| P6 | P6-3.1 | MGN-1_C07 | nd | 1 worker | SRX18307705 |
| P6 | P6-3.2 | MGN-1_D07 | *D. opacus/kohli* clade | 1 soldier | SRX18307705 |
| P6 | P6-4 | MGN-1_E07 | nd | 1 worker, 1 soldier | SRX18307705 |
| P6 | P6-reste | MGN-1_F07 | *D. opacus/kohli* clade | 4 workers, 4 soldiers | SRX18307705 |
| P6 | P6-1.1 | MGN-1_G08 | nd | 1 worker | SRX18307705 |
| P6 | P6-1.2 | MGN-1_H08 | nd | 1 soldier | SRX18307705 |
| P7 | P7-1.1 | MGN-1_G07 | nd | 1 worker | SRX18307705 |
| P7 | P7-1.2 | MGN-1_H07 | nd | 1 soldier | SRX18307705 |
| P7 | P7-2 | MGN-2_A10 | nd | 1 worker, 1 soldier | SRX18307706 |
| P7 | P7-3 | MGN-2_B10 | *D. sjoestedti/wilverthi* clade | 1 soldier | SRX18307706 |
| P7 | P7-4.1 | MGN-2_C10 | nd | 1 worker | SRX18307706 |
| P7 | P7-4.2 | MGN-2_D10 | nd | 1 worker | SRX18307706 |
| P7 | P7-4.3 | MGN-2_E10 | nd | 1 worker | SRX18307706 |
| P7 | P7-reste | MGN-2_F10 | nd | 2 workers, 3 soldiers | SRX18307706 |
| P8A | P8A-3.1 | MGN-2_A09 | nd | 1 worker | SRX18307706 |
| P8A | P8A-3.2 | MGN-2_B09 | nd | 1 soldier | SRX18307706 |
| P8A | P8A-4.1 | MGN-2_C09 | nd | 1 worker | SRX18307706 |
| P8A | P8A-4.2 | MGN-2_D09 | nd | 1 soldier | SRX18307706 |
| P8A | P8A-5.1 | MGN-2_E09 | nd | 1 worker | SRX18307706 |
| P8A | P8A-5.2 | MGN-2_F09 | nd | 1 soldier | SRX18307706 |
| P8A | P8A-reste | MGN-2_G09 | nd | 4 workers, 5 soldiers | SRX18307706 |
| P8A | P8A-2.1 | MGN-2_G10 | nd | 1 worker | SRX18307706 |
| P8A | P8A-2.2 | MGN-2_H10 | nd | 1 soldier | SRX18307706 |
| P9 | P9-1.2 | MGN-2_A08 | nd | 1 soldier | SRX18307706 |
| P9 | P9-2.1 | MGN-2_B08 | nd | 1 soldier | SRX18307706 |
| P9 | P9-2.2 | MGN-2_C08 | nd | 1 soldier | SRX18307706 |
| P9 | P9-3.1 | MGN-2_D08 | nd | 1 worker | SRX18307706 |
| P9 | P9-3.2 | MGN-2_E08 | nd | 1 soldier | SRX18307706 |
| P9 | P9-4.1 | MGN-2_F08 | nd | 1 worker | SRX18307706 |
| P9 | P9-4.2 | MGN-2_G08 | nd | 1 soldier | SRX18307706 |
| P9 | P9-reste | MGN-2_H08 | nd | 4 workers, 4 soldiers | SRX18307706 |
| P9 | P9-1.1 | MGN-2_H09 | nd | 1 worker | SRX18307706 |

**Supplementary Table 2:**
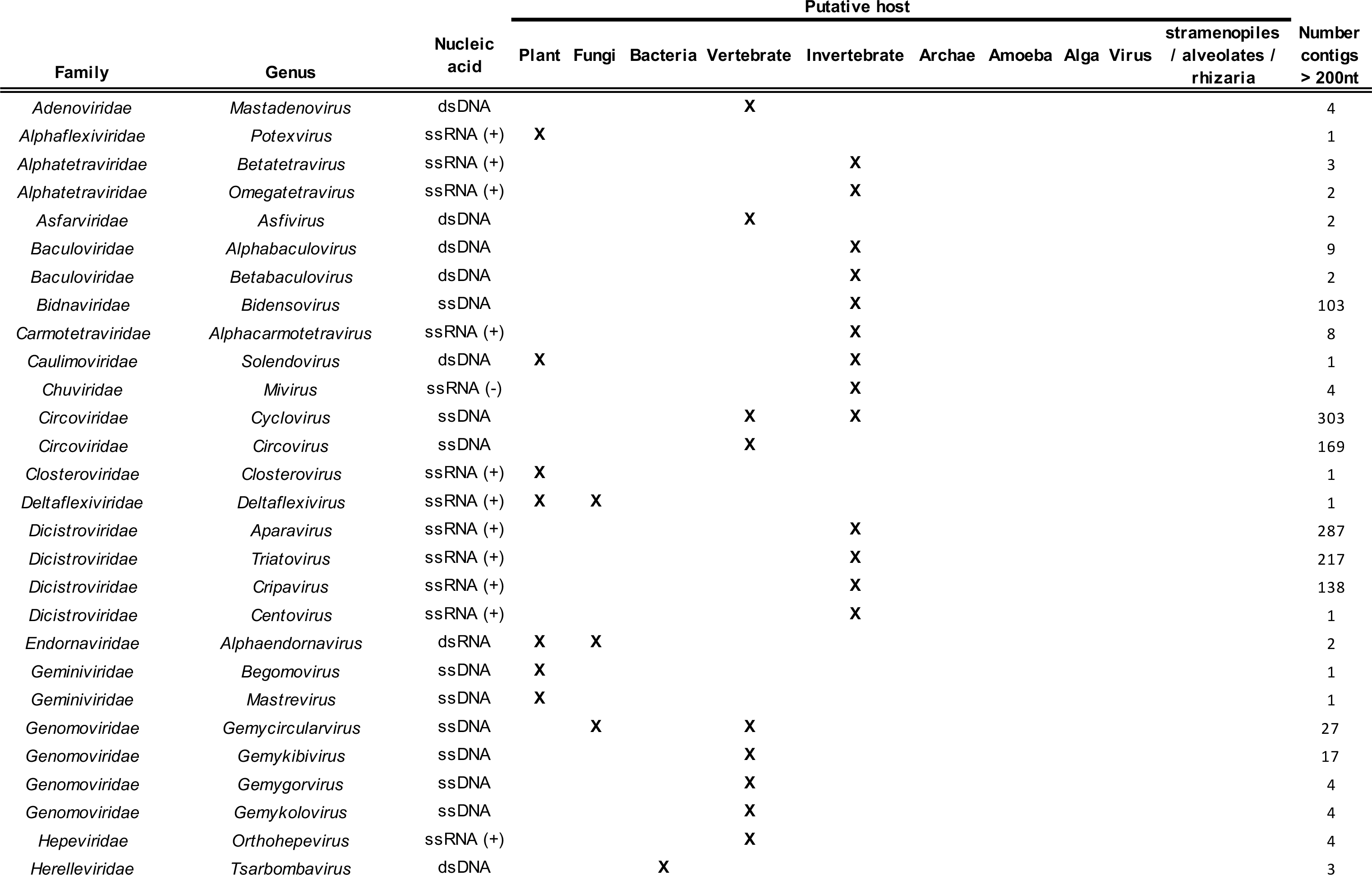
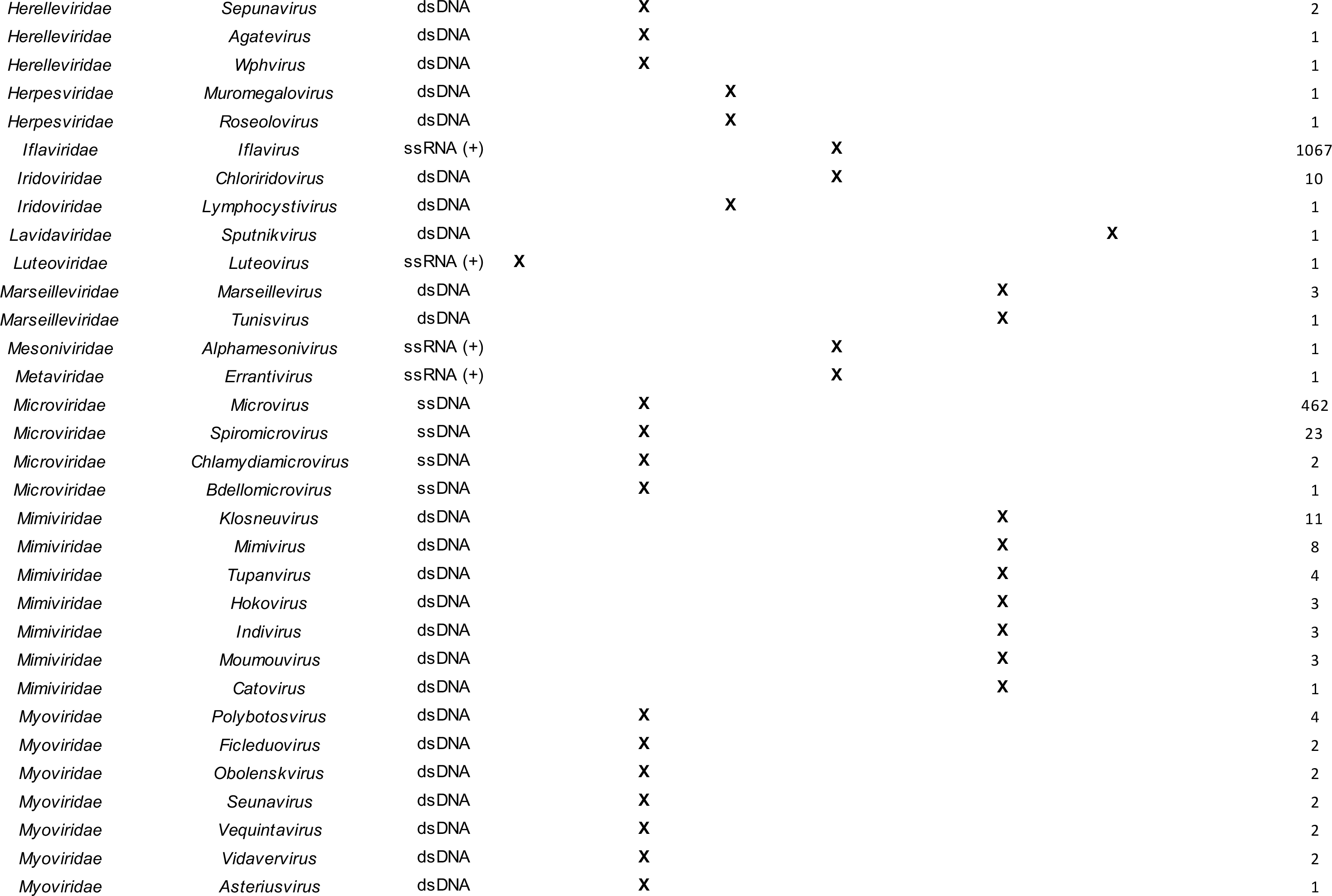
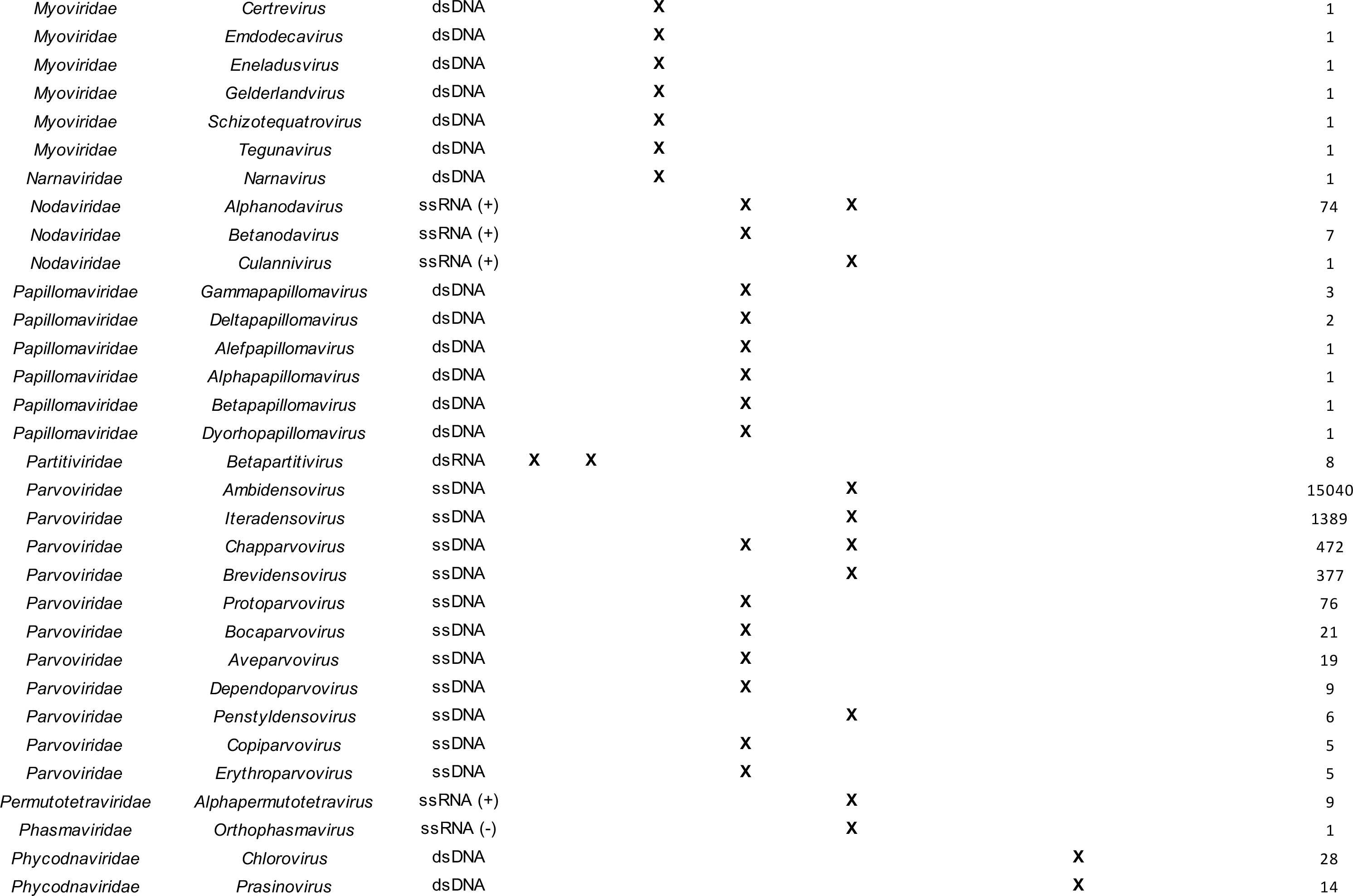
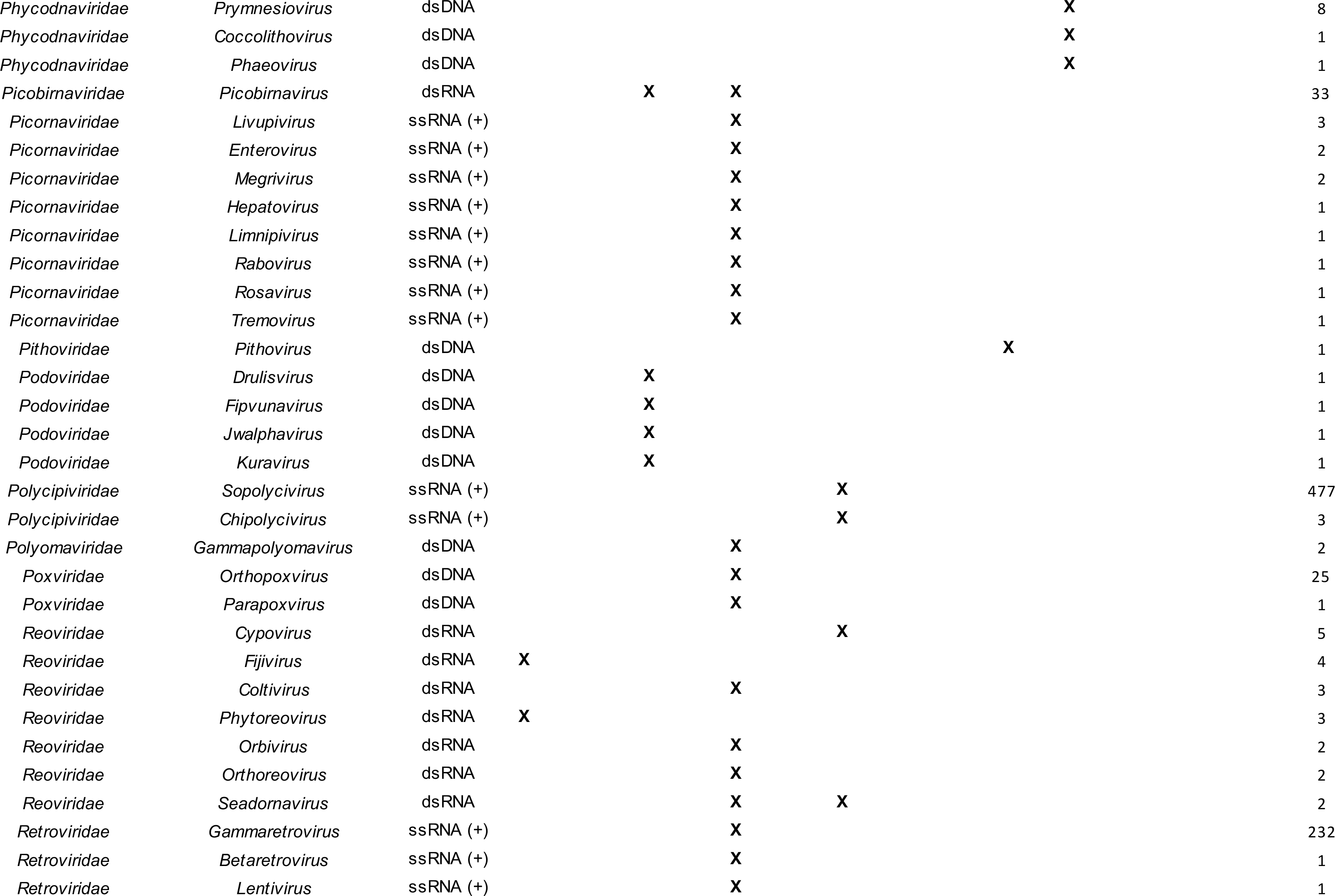
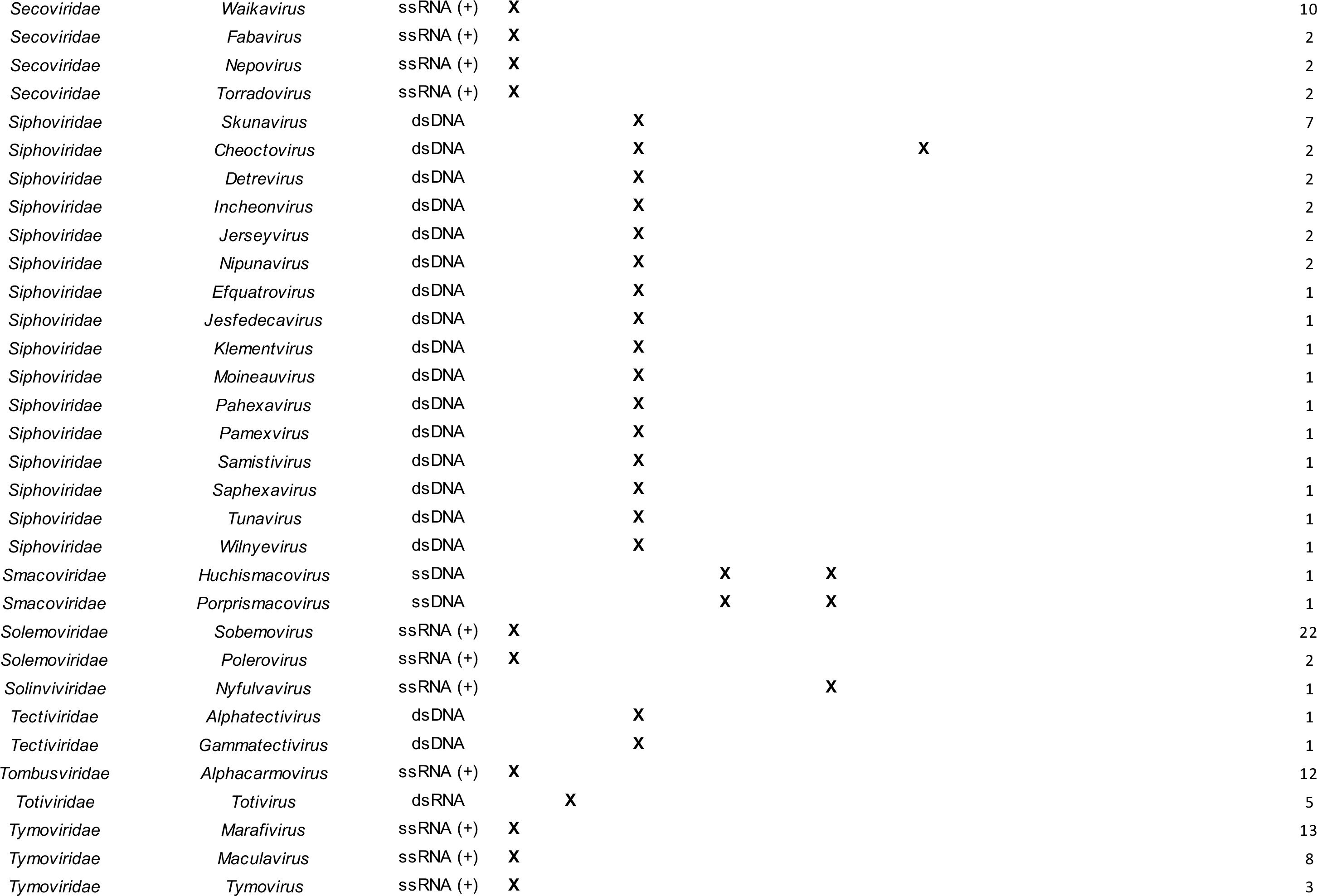
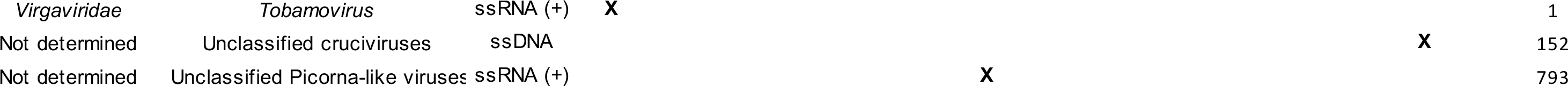
Number of virus-derived contigs yielding detectable homology to viruses using BLASTx (e-values < 0.001). Note that the *Myoviridae*, *Podoviridae* and *Siphoviridae* families were recently abolished.

**Supplementary Table 3:**
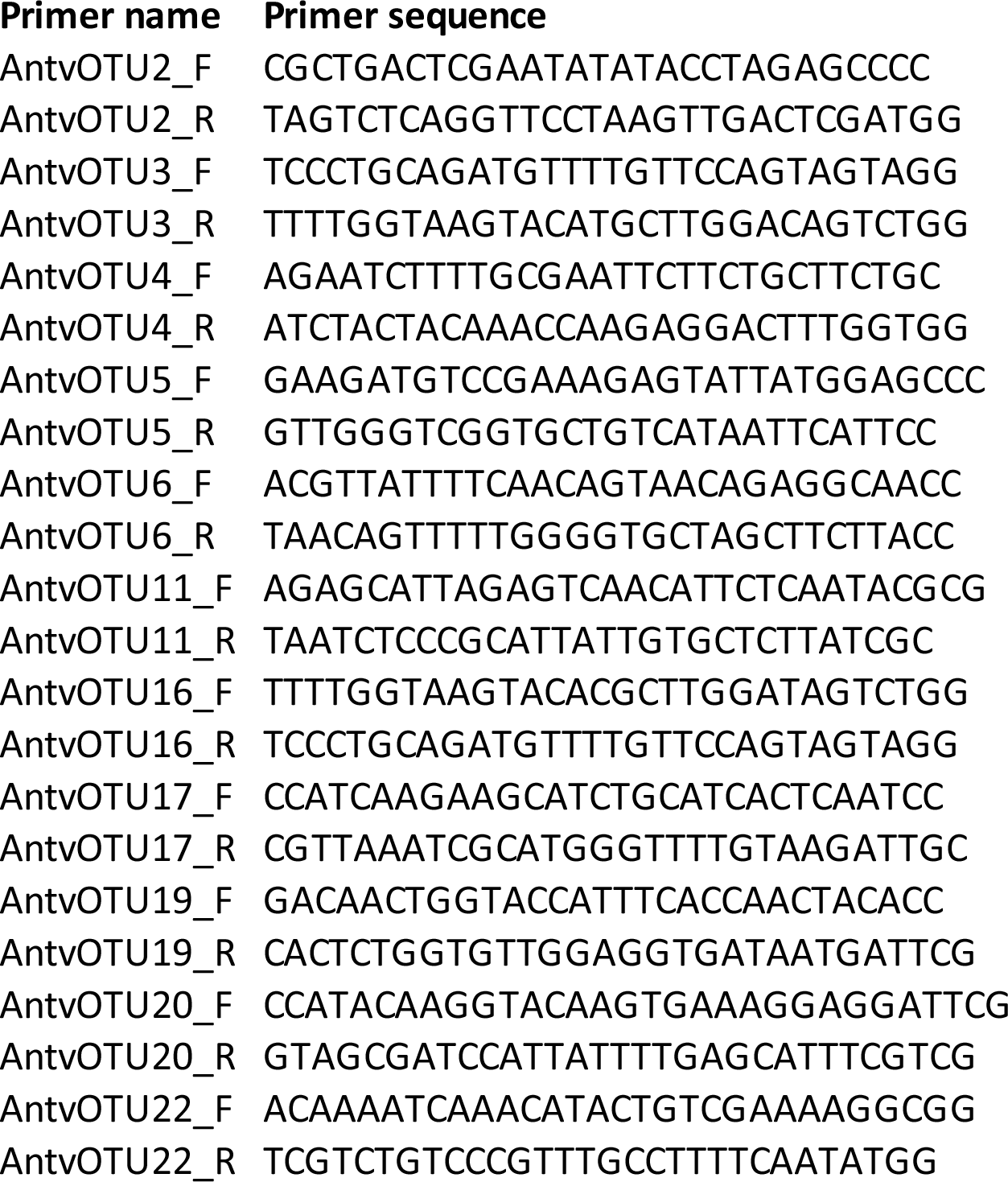
list of abutting primers used for amplifying the whole genomes of army ant associated cycloviruses

## Notes

https://doi.org/10.5281/zenodo.7636216

### Competing Interest Statement

The authors have declared no competing interest.

### Summary of Updates

This final version of the manuscript has been revised after having been reviewed by one Recommender and two reviewers of Peer Community in Infection.

